# Absolute copy number fitting from shallow whole genome sequencing data

**DOI:** 10.1101/2021.07.19.452658

**Authors:** Carolin M Sauer, Matthew D Eldridge, Maria Vias, James A Hall, Samantha Boyle, Geoff Macintyre, Thomas Bradley, Florian Markowetz, James D Brenton

## Abstract

Low-coverage or shallow whole genome sequencing (sWGS) approaches can efficiently detect somatic copy number aberrations (SCNAs) at low cost. This is clinically important for many cancers, in particular cancers with severe chromosomal instability (CIN) that frequently lack actionable point mutations and are characterised by poor disease outcome. Absolute copy number (ACN), measured in DNA copies per cancer cell, is required for meaningful comparisons between copy number states, but is challenging to estimate and in practice often requires manual curation. Using a total of 60 cancer cell lines, 148 patient-derived xenograft (PDX) and 142 clinical tissue samples, we evaluate the performance of available tools for obtaining ACN from sWGS. We provide a validated and refined tool called Rascal (<u>r</u>elative to <u>a</u>bsolute copy number <u>scal</u>ing) that provides improved fitting algorithms and enables interactive visualisation of copy number profiles. These approaches are highly applicable to both pre-clinical and translational research studies on SCNA-driven cancers and provide more robust ACN fits from sWGS data than currently available tools.

## Introduction

Somatic copy number alterations (SCNAs) are structural variants caused by ongoing chromosomal instability (CIN), and present as gain or loss of genomic regions that can include genes, regulatory elements, chromosome arms or whole chromosomes^1–4^. SCNAs affect a larger proportion of the genome than any other somatic alteration^5–8^, and are important drivers in many cancers^5,7,9,10^, particularly high CIN cancers, that typically lack actionable point mutations, are frequently associated with impaired DNA damage repair, and have poor treatment outcomes. Examples include high grade serous ovarian cancer (HGSOC)^11–13^, esophageal cancer^14,15^, head and neck cancer^16,17^, small cell lung cancer^18,19^, lymphoma^20^, neuroblastoma^21^, and triple negative breast cancer^22,23^. Reliable characterisation of SCNAs in the clinic can improve early detection of cancer^24–26^, and will be critical for molecular stratification of patients with high CIN cancers for targeted cancer therapies^27,28^.

Despite their importance, it remains challenging to reliably identify and quantify SCNAs in commonly used laboratory cancer models and clinical samples. Massively parallel whole genome sequencing (WGS) based approaches have replaced older array-based methods to detect and estimate DNA copy number genome-wide^29–31^. However, these approaches usually require high sequencing depth associated with significant costs, or are highly sensitive to DNA quality, making them unsuitable for formalin-fixed paraffin-embedded (FFPE) tissues^32^, which are the most commonly available diagnostic samples.

In contrast, low coverage or shallow whole genome sequencing (sWGS) offers a cost-effective alternative to detect SCNAs in various sample types, including FFPE samples^33–35^. Several read depth-based bioinformatic tools, such as QDNAseq^35^, have been developed to facilitate the detection of copy number changes from sWGS^36^, and outperform array-based methods^37^. sWGS applications also have significantly lower computational and data storage requirements in comparison to high coverage sequencing. Consequently, sWGS is now widely accessible in both clinical and research settings.

The commonest representation of SCNAs is on a relative scale, in units of normalised read counts for a given genomic region relative to the median normalised read counts across the whole genome of a sample. However, relative measures of SCNAs are difficult to interpret because they do not provide direct information on a sample’s ploidy, are highly dependent on the tumour purity (the proportion of normal contaminating cells within a tumour sample) and may be further confounded by intra-tumour heterogeneity. This means that SCNA estimates from samples in the same cohort, or even the same patient, are not directly comparable. Instead, SCNAs measured on an absolute scale (in ‘copies per cancer cell’) are required for accurate interpretation of copy number changes^30^ and inter-sample comparisons. Interpreting relative SCNA as absolute SCNA is a common mistake and results deriving from this misinterpretation overrepresent differences arising from tumour purities and ploidies rather than biological differences in the copy number landscape across different samples. While several deep WGS-based tools are already available to determine ACN, the capability to reliably assay absolute SCNAs from sWGS data is much less developed, making this analysis difficult for non-experts. This deficiency frequently leads to deep WGS-based tools being used in a “black box” manner, leading to questionable results^38^.

Here, we provide an overview of the underlying principles for determining absolute copy number (ACN) from sWGS and demonstrate the performance of currently available copy number tools for typical laboratory and translational experiments. We explore current challenges of estimating ACN by focussing on the three main confounding factors: 1) the tumour purity^39^, 2) the ploidy^40^ or average DNA content of a tumour, and 3) the extent of heterogeneous tumour cell populations from clonal evolution^41^. Understanding these well-established concepts and their involvement in determining copy number signal is critical to enable comparative and robust SCNA interpretation from cost-effective assays, such as sWGS. To make ACN data analyses widely accessible to the research community, we provide detailed insights into workflows and propose the use of an improved and validated tool, called Rascal (<u>r</u>elative to <u>a</u>bsolute copy number <u>sca</u>ling), which enables more robust ACN fitting from sWGS. Rascal is available as an R package as well as an interactive web application to visualise ACN data, choose between competing ACN solutions, and manually curate difficult-to-fit samples.

## Results

### Evaluation of absolute copy number fitting tools on high purity samples

Absolute copy number fitting from whole genome sequencing requires modelling of genomic copy number data as a function of the cell ploidy and tumour purity from the sample of interest (see Methods). The relative copy number obtained for any region of the genome will always reflect combined contributions from the tumour and normal contaminating diploid cells, depending on the sample’s tumour purity. For example, in a triploid tumour, a cellularity of 0.7 (i.e. 70% of cells are cancerous and 30% are normal) will lead to an observed copy number of 0.7 × 3 + 0.3 × 2 = 2.7. Thus, the major impact of contaminating normal cells is to reduce the copy number signal from cancer cells, resulting in lower gains and losses (Supplementary Fig. 1 and Supplementary Video 1). Several bioinformatic tools estimate purity and ploidy to facilitate ACN fitting (Extended Data Table 1 provides an overview and description of tools). Most tools, including TITAN^42^, and ASCAT^43,44^, were developed for the analysis of deep WGS/SNP array data, estimating allele-specific ACN with a focus on determining tumour heterogeneity and loss of heterozygosity. In addition to requiring high coverage sequencing data, many of these tools also rely on sequencing reads from matched normal samples. Matched normal or germline DNA, however, is often difficult to obtain for laboratory and clinical samples, and is not available for the vast majority of cancer cell lines. ACE^45^ is, to our knowledge, the only tool that has been specifically designed for sWGS data and does not require input from matched normal sequencing reads. However, ichorCNA^25^ and ABSOLUTE^46^, have also been applied to various sWGS-based studies, although both recommend the use of matched normal samples.

**Figure 1.**
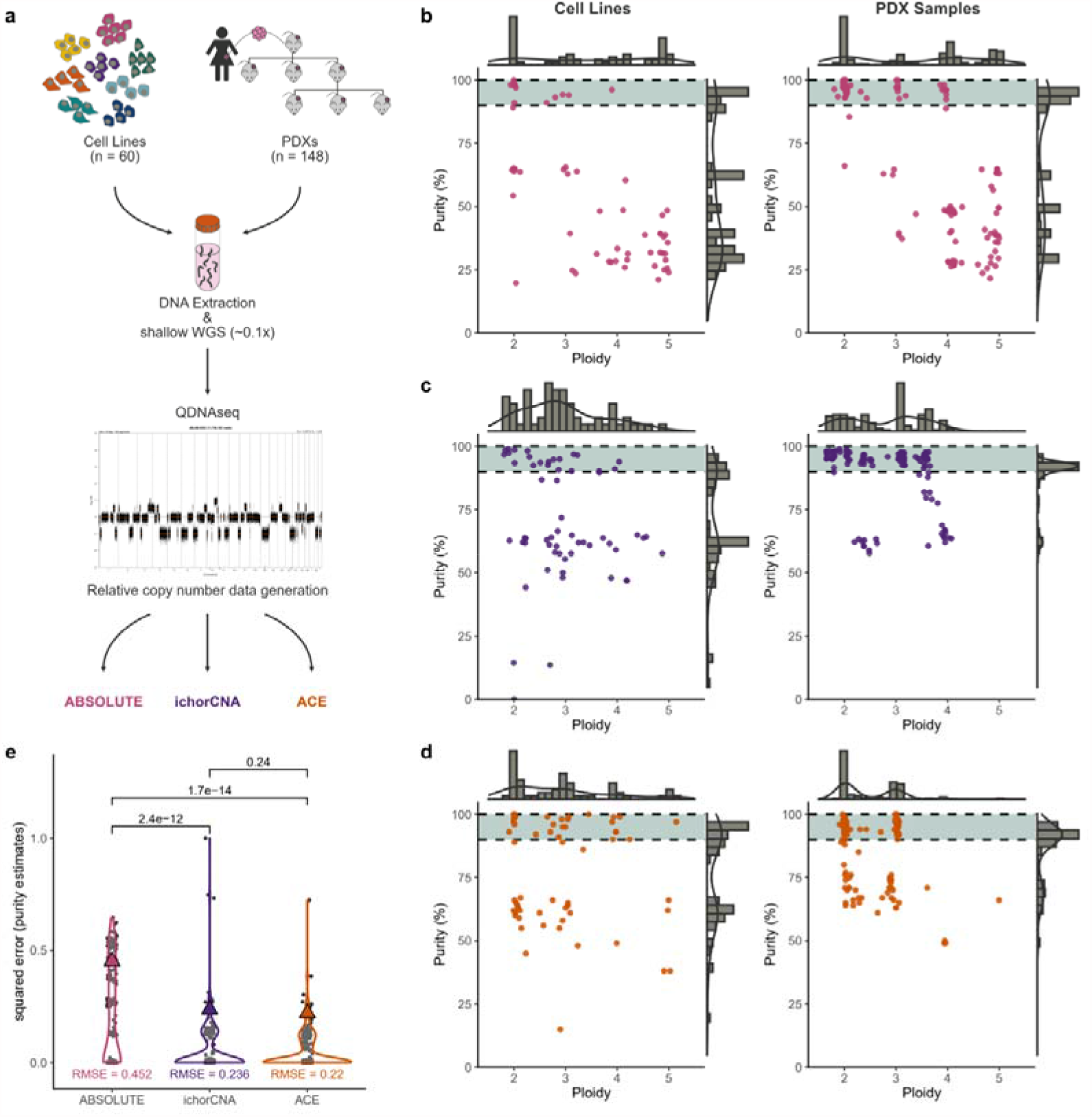
Performance evaluation of current ACN fitting tools. (**a**) Study design for absolute copy number (ACN) tool performance testing. High purity samples comprising a total of 60 cell lines and 148 PDX samples generated from 14 patients were subjected to sWGS. Relative copy number data was generated using QDNAseq, and subsequently analysed with ABSOLUTE, ichorCNA and ACE to test ACN fitting performance. Scatter plots of ploidy and purity estimates are shown for ABSOLUTE (**b**), ichorCNA (**c**) and ACE (**d**). The expected purity estimates of 90-100% are indicated by the green-shaded areas. Summary distributions of purity and ploidy estimates are shown on the margins of each plot. (**e**) Comparison of ABSOLUTE, ichorCNA and ACE using the squared errors. Assuming purities of 100%, P-values were calculated on the squared errors using the paired one-sided Wilcoxon test. RMSE (root mean squared error) values are indicated by coloured triangles.

To objectively evaluate the performance of ABSOLUTE, ichorCNA and ACE in common pre-clinical research settings, we used sWGS data from a total of 60 ovarian cancer cell lines and 148 PDX samples derived from 14 individual patients (Fig 1a). These sample types have high tumour purity, as confirmed by inspection of *TP53* mutant allele fractions (MAFs) from Tagged-Amplicon sequencing^47^ (TAmSeq) data (Supplementary Fig. 2), and therefore provide a simple setting for ACN fitting as only two confounding factors (ploidy and subclonality, but not purity) have to be considered. sWGS data from these datasets were analysed using QDNAseq to generate read count and segmented relative copy number data. ABSOLUTE, ichorCNA and ACE were then applied to estimate sample purity and ploidy. The best or highest ranked solution for each sample from each tool was taken forward for further analysis. Only 28–53% of cell lines and 49–82% of PDX samples were correctly estimated to have a tumour purity of ≥ 90%, indicating a common bias for all three tools to significantly underestimate tumour purity (Fig. 1b-d). ACE and ichorCNA both performed better than ABSOLUTE (P = 1.7 × 10^−14^ and P = 2.4 × 10^−12^, respectively, paired one-sided Wilcoxon test comparing squared errors), with ACE having the lowest root mean square error (RMSE) of 0.22, followed by RMSE = 0.24 for ichorCNA and RMSE = 0.45 for ABSOLUTE (Fig. 1e). This shows that current unsupervised fitting algorithms do not consistently yield correct ACN solutions from sWGS data.

**Figure 2.**
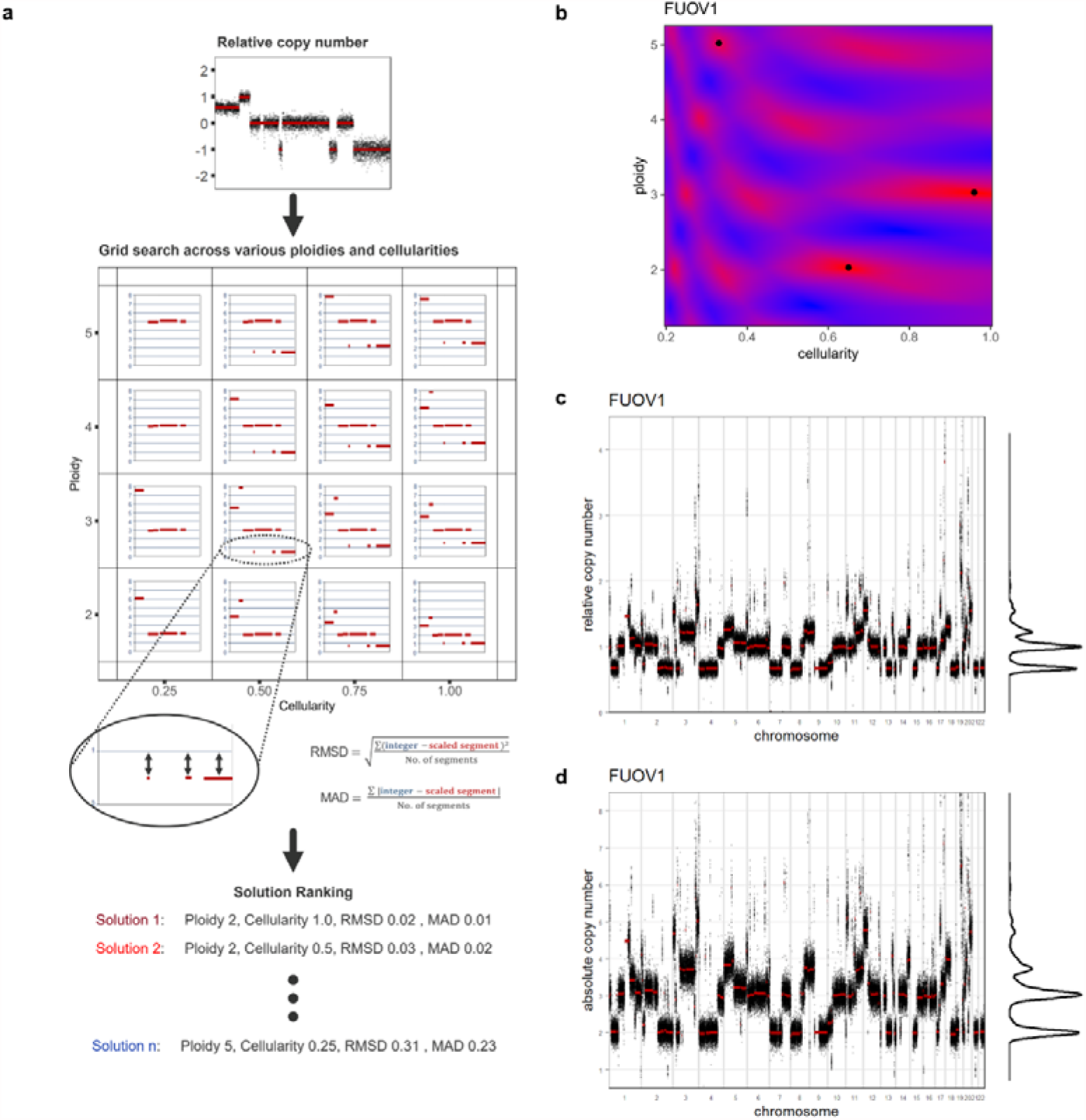
Underlying principles and ACN fitting procedure using Rascal. (**a**) Underlying principles of Rascal. Segmented relative copy number (RCN) data (from QDNASeq) is input into Rascal and a grid search is subsequently performed to fit the RCN data to various ploidies and cellularities. Rascal assesses the goodness-of-fit for each ploidy-cellularity solution using a mean absolute deviation (MAD) or root mean square deviation (RMSD) distance function by estimating the absolute difference between the scaled (fitted) copy number values (red segments) and the nearest whole copy number integer (blue horizontal lines) for each segment. The MAD or RMSD distance measures are compared across various model fits to rank all ACN solutions for each sample and to determine the best fit (lowest distance function). An example using the FUOV1 cell line is shown in (**b**-**d**): (**b**) Heatmap representing the grid search applied by Rascal, with cold areas (poor fits) shown in blue, and hot areas (good fits) shown in red. The best solutions (local minima) are indicated as black dots. Out of these, the solution with the lowest MAD distance has a ploidy of 3.03 and cellularity of 0.96. The solution with the highest MAD distance has a ploidy of 5.02 and a cellularity of 0.33. (**c**) Relative copy number plot and (**d**) fitted absolute copy number plot (best solution – ploidy 3.03 and cellularity 0.96) for FUOV1 with density distributions of copy number segments shown to the right of each plot.

### Rascal – an improved method for accurate ACN fitting from sWGS data

We next investigated why current tools frequently result in suboptimal fits, focussing on ACE. Our observations show that ACE underestimates tumour purity, particularly for samples with noisy copy number profiles. This bias occurs because ACE’s “goodness-of-fit” function computes differences between segmented copy numbers on the relative scale, where decreasing cellularity leads to more closely spaced copy number steps and thus smaller distances (Supplementary Fig. 1 and Supplementary Video 1). ACE provides customizable penalty factors which could compensate for low cellularity solutions and ploidies that diverge from the diploid state, but the ploidy factor is unsuitable for cancer types where whole genome duplication is frequent. Additionally, ACE restricts its solution space for copy number states to values ≤ 12, limiting its applicability to high CIN cancers—for example MYC amplifications of > 20 copies are frequently observed across many cancer types^3,5,7^.

Based on these observations, we propose an improved ACN fitting procedure, called Rascal (<u>r</u>elative to <u>a</u>bsolute copy number <u>sca</u>ling), that builds on ACE and leads to more plausible solutions, especially for poorer quality copy number profiles. The full mathematical framework of Rascal is provided in the Methods section. Fig. 2a summarises the main principles of Rascal. Like ACE, Rascal considers various combinations of a range of different ploidy and cellularity values using a grid search algorithm (Fig. 2a). However, Rascal estimates the “goodness-of-fit” function in the absolute, not relative, space. The distance function is calculated either as the root mean square deviation (RMSD) or mean absolute deviation (MAD) and provides a measure that can be compared across various model fits to determine the best fit from the lowest distance for each sample (the choice of distance function is discussed in the next section). Fig. 2b provides an example using the triploid ovarian cancer cell line, FUOV1^48,49^. Using the MAD fitness measures, Rascal finds that the best solution (lowest MAD) has a ploidy of 3.03 and cellularity of 0.96 (Fig. 2c, d).

To further validate ploidy and cellularity predictions from Rascal, we applied it to our high purity cell line and PDX datasets. Expected fits of ≥ 90% cellularity were achieved for 93% and 94% of cell line and PDX samples, respectively (Fig. 3a). ACN fits were not achieved for three cell lines and two PDX samples. To test the accuracy of these predictions, we compared fitted ploidy estimates to previously published ploidy data for 30 ovarian cancer cell lines from the Cancer Cell Line Encyclopaedia^48^, the Sanger Cell Model Passports^49^, or obtained from chromosome banding and metaphase spreads^50–54^ (Extended Data Fig. 1a-d; Supplementary Table 1 and 2). Notably, external data showed discordant ploidy estimates for four out of 12 cell lines for which multiple sources/methods were available (CAOV3, CAOV4, OV56, and IGROV1).

**Figure 3.**
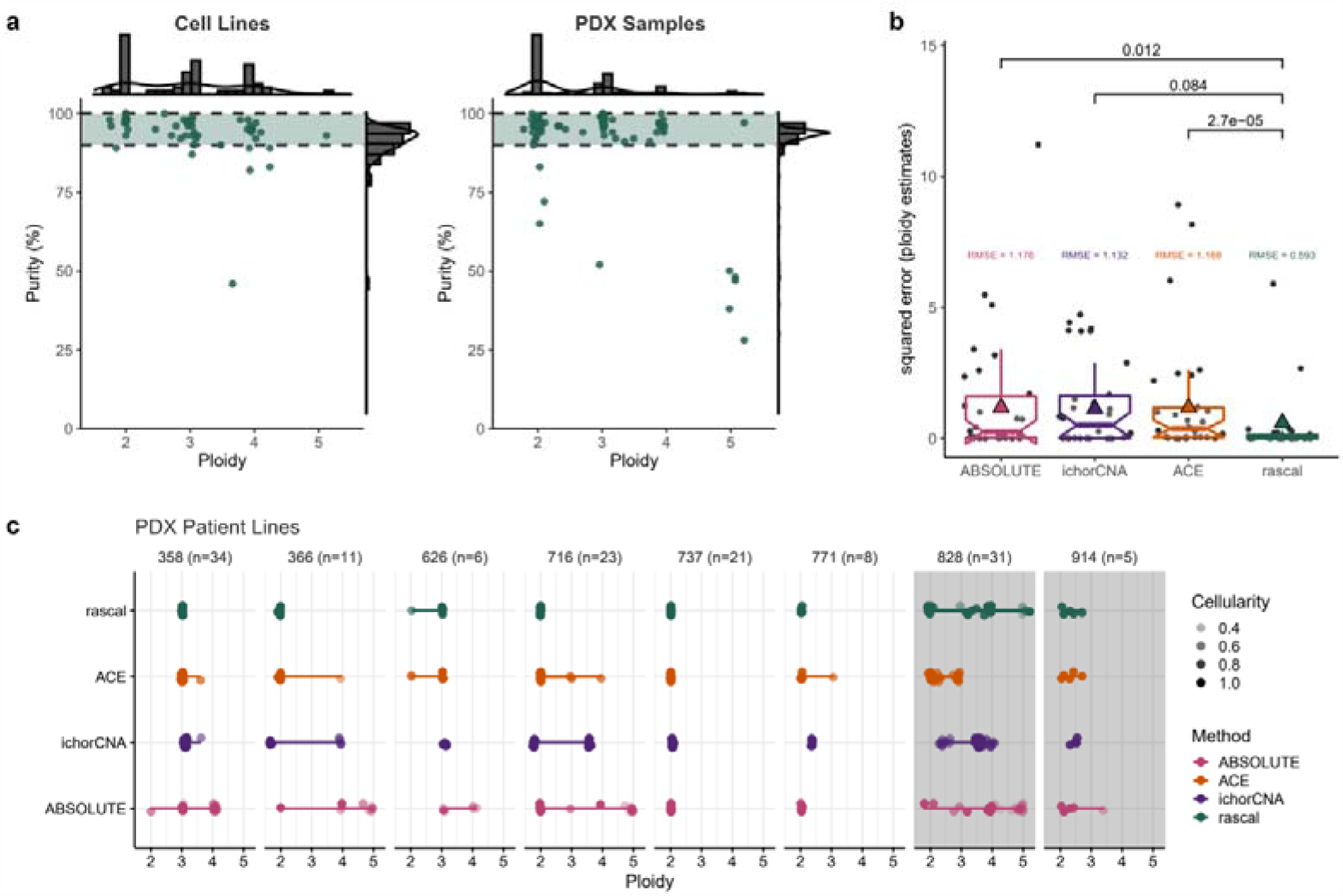
Rascal method validation and benchmarking on high purity samples. (**a**) Rascal fits (ploidy and purity estimates) for cell line (left) and PDX (right) samples. Expected target area (90-100%) for purity estimates is indicated by a green-shaded ribbon, and distributions for both purity and ploidy estimates are shown on the margins of each plot. (**b**) Comparison of ABSOLUTE, ichorCNA, ACE and Rascal using previously published ploidy cell lines data (also see Extended Data Fig.1 and Supplementary Table 1 and 2). Data is shown as squared errors comparing fitted ploidy estimates to previously reported ploidies. P-values were calculated on the squared errors using the paired one-sided Wilcoxon test. RMSE (root mean squared error) values are indicated by coloured triangles. (**c**) Ploidy comparison across PDX samples from the same patient line for the four different ACN tools (ABSOLUTE, ichorCNA, ACE and Rascal). Associated cellularities are indicated by the opacity of the data points. Chromosomal unstable PDX lines (828 and 914) are shaded in grey (also see Supplementary Video 2 for PDX line 914 and Supplementary Video 3 for ACE fits for PDX line 716).

As shown in Extended Data Fig. 1, ploidy fits obtained using Rascal are consistent with previously published ploidy measures (< 0.5 copy number steps difference), with the exception of OVCAR-5 and OV4453. For the four cell lines with conflicting published ploidy measures, Rascal’s ploidy estimates were concordant with at least one of these measures. In comparison, ACE, ichorCNA and ABSOLUTE only achieved ploidy estimates matching those previously published in ≤ 50% of cell line samples. For ploidy estimates, Rascal performed significantly better than ACE and ABSOLUTE (P = 2.7 × 10^−5^ and P = 0.012, respectively; paired one-sided Wilcoxon test) and had the lowest RMSE value of 0.59, followed by ichorCNA (RMSE = 1.13), ACE (RMSE = 1.17) and ABSOLUTE (RMSE = 1.176) (Fig. 3b).

Copy number profiling of 1,451 samples from 509 PDX models showed that SCNAs and sample ploidies are highly conserved between patients and different PDX generations, and from multiregional sampling. Consistent with this data, Rascal, but not ACE, ichorCNA or ABSOLUTE, resulted in highly consistent fits for different generations of PDX samples derived from the same patient, with the exception of one mis-fitted sample (as indicated by the low cellularity estimate) in patient line 626 (Fig. 3c). PDX samples from patient line 828 and 914 showed ongoing CIN with changes in SCNA between different PDX animals and generations. For example, in PDX line 914 there was shifting of segments from chromosome 6 and 7 by ≥ 1 copy number steps resulting in different sample ploidies, with the remainder of the genome remaining stable across different PDX animals (Supplementary Video 2). In comparison, ACN profiles for PDX line 716 obtained using ACE showed shifting of copy number segments across the whole genome by ≥ 1 copy number steps for samples with differing ploidy states indicating incorrect ACN fits (Supplementary Video 3).

### MAD is the preferred distance function for accurate ACN fitting

Although accurate estimation of sample purity and ploidy are essential for ACN fitting, a sample’s heterogeneity or subclonality is the third critical factor that has to be considered. The main assumption made during ACN fitting is that all tumour cells within a sample have the same copy number states, i.e. the tumour is homogeneous. In reality, tumours may display considerable heterogeneity with complex clonal architectures. Unique gains and losses of genomic regions in tumour subclones will all contribute to the average copy number profile, resulting in copy number segments that cannot be accurately scaled without further information provided by deeper sequencing and therefore appear as segments falling between integer states. Since the “goodness-of-fit” for each ACN solution is estimated based on how close a scaled copy number segment aligns to its nearest full integer state, we compared the performance of the two error functions (RMSD and MAD) across our cell lines and PDX samples.

RMSD and MAD distance functions resulted in concordant solutions for the majority of PDX and cell line samples (86% for cell line and 87% for PDX samples; Fig. 4a). Examining the discordant models (red; Fig 4a), the MAD distance function fitted 24 out of 27 (88.9%) to cellularities ≥ 90% (Fig. 4b). In contrast, using the RMSD distance function, only four (14.8%) of the discordant samples were fitted to cellularities ≥ 90%, suggesting that the RMSD distance function is more prone to underestimating cellularity in these cases. We also compared RMSD- and MAD-fitted ploidies to published ploidy data which was available for five out of the eight discordant cell line samples (Fig. 4c). Interestingly, these included three out of the four cell lines (CAOV3, CAOV4 and OV56) for which varying ploidy values had previously been reported (Supplementary Table 2). While the MAD distance function found matching ploidy solutions for all five cell line samples, the RMSD distance function was highly dissimilar to published data, with the exception of CAOV3 and CAOV4 (Fig. 4c). To determine the correct ploidy, we prepared metaphase spreads for CAOV3 and CAOV4 (Fig. 4d) which confirmed accurate ploidy fits from the MAD distance function.

**Figure 4.**
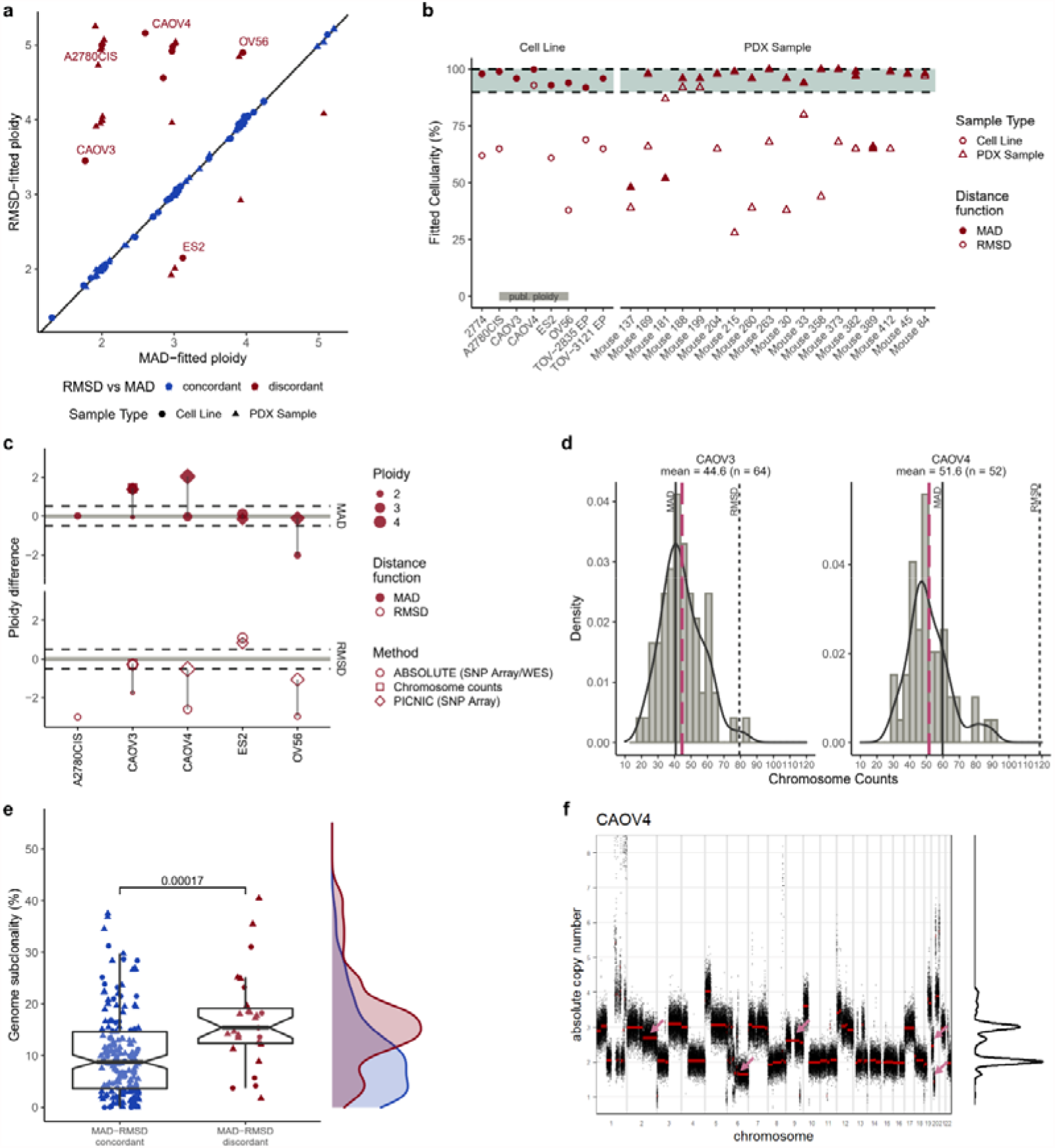
RMSD vs MAD goodness-of-fit function and tumour subclonality. (**a**) Comparison of fitted ploidy estimates using the RMSD or MAD distance function for cell lines (circles) and PDX samples (triangles). Samples with concordant ploidy fits for both RMSD and MAD are shown in blue, discordant fits are shown in red. Discordantly fitted cell lines for which published ploidy data was available (see Extended Data Fig.1 and Supplementary Table 2) are highlighted by labels. (**b**) Comparison of MAD and RMSD fitted cellularities for all discordant (red) samples from (a). Cell line samples are shown as circles, PDX samples as triangles. Cellularity fits obtained from the RMSD function are shown as hollow objects, whereas MAD-fitted cellularities are indicated as filled object. Expected cellularity estimates of 90-100% are indicated by the shaded area. (**c**) Bland-Altman plot comparing MAD (filled) and RMSD (hollow) fitted ploidies to published ploidies available for five cell lines. The y axis shows the difference between published and fitted ploidies. Ploidy difference of +/−0.5 copy number steps is indicated by dashed lines. Shapes indicate methods for determining published ploidies. For two cell lines (CAOV3, and CAOV4) both distance functions produced plausible but different fits. To confirm which solution is correct, metaphase spreads were performed on CAOV3 and CAOV4 to count the number of chromosomes present in both cell lines (**d**). Mean chromosome counts are indicated by dashed pink lines. Expected chromosome counts for MAD and RMSD ploidy estimates are indicated by a solid or dotted grey line, respectively. (**e**) Percentage of genome subclonality calculated as the total length of segments at intermediate copy number states divided by the total length of all segments (i.e. length of the genome) for MAD-RMSD concordant, and discordant samples. The genome subclonality density plot for each of the two groups is shown to the right. P-values were calculated using the unpaired one-sided Wilcoxon test. (**f**) Fitted absolute copy number profile for CAOV4 using the top ranked MAD solution (ploidy 2.6; cellularity 1). Intermediate/subclonal copy number segments are indicated by pink arrows.

To understand whether the RMSD-MAD discrepancy is caused by the presence of subclonal states, we estimated the fraction of subclonal segments (genomic subclonality) as the total length of segments at subclonal/intermediate copy number states (≥ 0.25 copy number steps distance to the nearest integer) divided by the total length of the genome. Samples with RMSD-MAD discordant fits had significantly higher proportions of genomic subclonality compared to samples for which RMSD and MAD achieved concordant fits (P= 0.00017, Wilcoxon test; Fig. 4e). This is illustrated using CAOV4 as an example (Fig. 4f). The correct solution has a ploidy of 2.6 and cellularity close to 1 and has the lowest MAD. The absolute copy number profile shows the presence of subclonal/intermediate segments on chromosome 2, 6, 9 and 20. In contrast, the RMSD function favours a set of three solutions with similar distance measures containing ploidies of 5.16, 3.17 and 4.17 (Supplementary Fig. 3a). Of these, only the ploidy 5.16 solution has a cellularity estimate close to 1 but appears to “overcompensate” for the segments on chromosomes 2, 6, 9 and 20 (Supplementary Fig. 3b), resulting in a higher ploidy fit.

To further explore the effect of subclonal populations on ACN fits, we provide *in silico* models of subclonality in Extended Data Fig. 2 (see also Supplementary Fig. 4, Supplementary Video 2). These indicate that Rascal can obtain accurate fits providing the dominant clone comprises >55–75% of the sample tumour content. In addition, our genome subclonality estimate (Fig. 4e) strongly correlated with the *in silico* simulated subclonal fraction (R = 0.69 for A2780 mixtures, and R = 0.85 for PDX line 914 mixtures; Pearson Correlation) (Extended Data Fig. 2d).

While both distance functions produce equivalent ACN fits for the majority of samples, we show that the MAD function produces more accurate fits in cases where subclonal states might influence parts of the copy number profile. The MAD function thus produces better purity and ploidy estimates and reveals intermediate copy number values as an indicator of tumour heterogeneity.

### Clinical tissue samples require prior knowledge to guide accurate ACN fitting

Cancer specimens from patients present the most complex challenges for ACN fitting because these samples have highly variable purities with unknown ploidies. These limitations make it hard to choose between multiple predictions for best fits when differing ploidies and cellularities result in the same optimum distance measure (see Methods). This phenomenon of competing best fits was observed in 52% of cell line, 35% of PDX samples, and 36% of clinical tissue samples (Supplementary Fig. 5a). In pure samples, competing best fits can be easily distinguished using the assumption of high purity (Supplementary Fig. 5b, c). For clinical tissues, other prior knowledge regarding ploidy or cellularity is therefore ideally required to guide accurate ACN fitting. This additional knowledge could be obtained for example through flow cytometry, fluorescent *in* situ hybridisation or computational histopathology approaches. Another important surrogate for tumour cellularity is the mutant allele fractions (MAFs) for clonal driver mutations, such as *APC* or *KRAS* in colorectal cancer^6,55^ and *PIK3CA* in ∼50% of breast cancers^6,56^.

**Figure 5.**
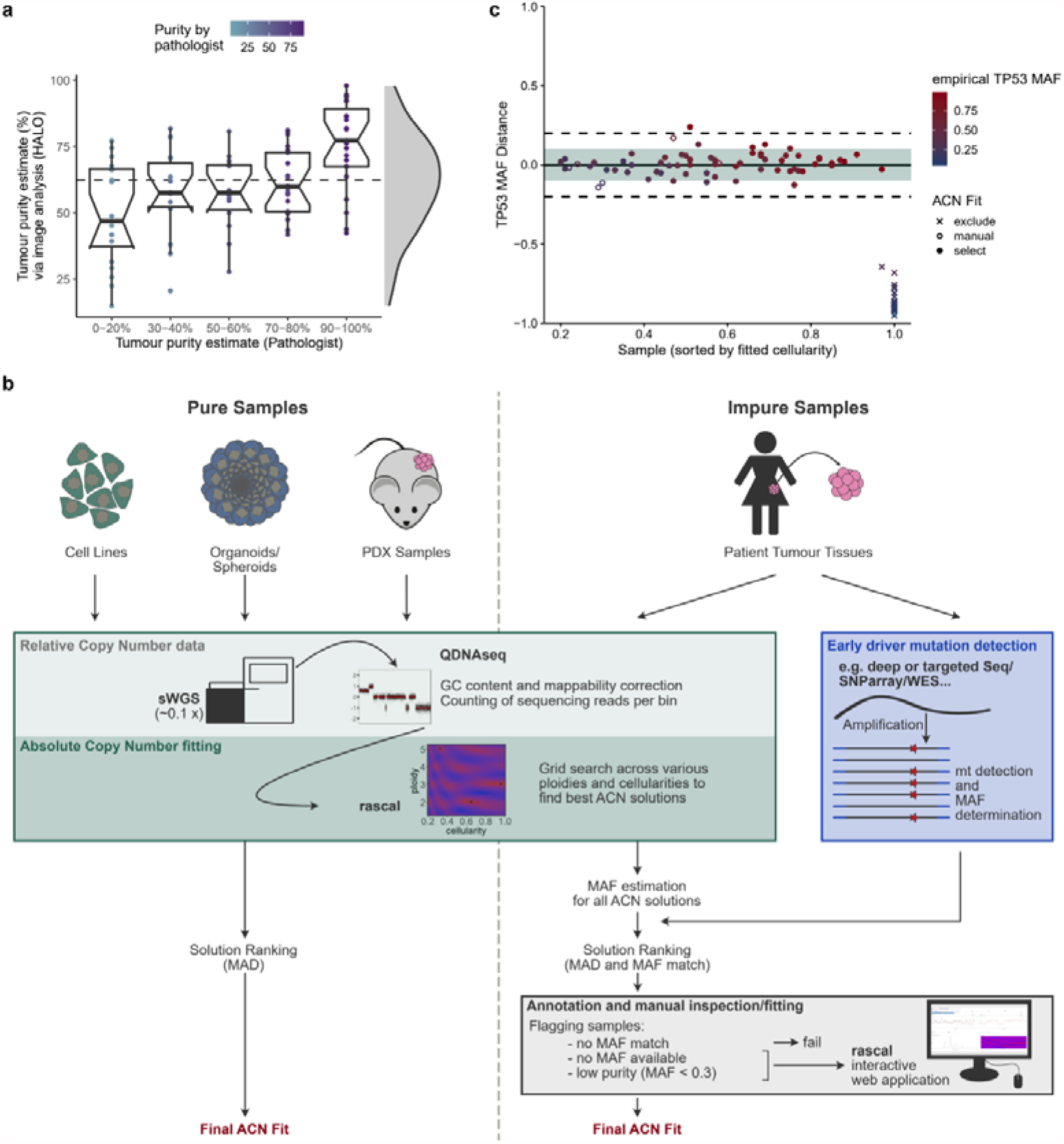
Competing best fit solutions and fitting of clinical tissue samples. (**a**) Purity estimates for 97 HGSOC FFPE tissue samples. Samples are shown grouped according to (underestimated) histological purity estimation performed by a pathologist, compared to histological purity estimates via computational image analysis (HALO software). The distribution of HALO-estimated purities is shown on the right, with a median purity of 67.7% (indicated by grey dashed line). (**b**) Adapted workflow to allow accurate ACN fitting of clinical (impure) samples in comparison to the basic workflow for pure samples (e.g. cell lines, organoids/spheroids and PDX samples). In short, samples are subjected to sWGS, and segmented relative copy number (RCN) data is generated using QDNASeq. Rascal subsequently performs a grid search on the RCN data across various ploidies and cellularites to find putative ACN solutions. For impure samples expected mutant allele fractions (MAFs) are estimated for early driver genes, e.g. *TP53*, for each ACN solution and compared to empirical MAFs obtained from deeper sequencing or SNP array data. Solutions are ranked based on the MAD distance function and comparison of estimated to empirical MAF values. Samples for which no suitable ACN solution is found (based on MAF comparison) are excluded from downstream analyses. Samples for which no empirical MAF values are available or are of very pure purity (< 0.3 MAF) are flagged for manual inspection to either confirm or correct ACN fits using the interactive web application of Rascal allowing manual inspection of sample fits. (**c**) Bland-Altman plot showing the difference between empirical and estimated *TP53* MAFs for selected ACN solutions across our clinical HGSOC cohort (n=132). Samples are sorted by their fitted cellularities, and colour-coded by their empirical *TP53* MAF values. Samples that were manually fitted using the interactive web application are indicated by hollow circles (n=6), whereas samples for which no ACN fit could be obtained and which were consequently excluded are shown as crosses (n=30). Note that all of these samples were fitted to ploidy 2/cellularity 1 solutions, owing to the high fraction of normal contaminating cells as indicated by low *TP53* MAFs (median 0.118; IQR 0.093).

We recommend a workflow using Rascal for clinical tissue samples (Fig. 5b) which facilitates the incorporation of prior tumour purity or cellularity knowledge as well as visual inspection of “difficult-to-fit” samples using the Rascal’s interactive web application interface (see Methods).

To illustrate this, we used a total of 142 clinical tissue samples, encompassing 23 normal fallopian tube (FT) and 119 HGSOC samples, which ranged from 15–98% cellularity estimated by a clinical pathologist (Fig. 5a). Since *TP53* mutation is a ubiquitous early driver event in HGSOC, we used *TP53* MAFs as a surrogate for cellularity. Rascal was able to generate possible ACN solutions for 132 out of the 142 (93%) samples. Out of these, 30 samples failed ACN fitting and were excluded owing to low *TP53* MAFs (median 0.12; IQR 0.09) suggesting low purity. Out of the remaining 102 samples, six were identified as “difficult-to-fit” and reviewed in detail using the visual Rascal interactive web application and manually re-fitted. Fig. 5c shows the comparison of fit-based estimated *TP53* MAFs and *TP53* MAFs from sequencing data.

To test the performance of Rascal in lower cellularity samples, we generated *in silico* mixtures of tumour:normal DNA using sequencing reads from a high purity tumour sample and a matched normal fallopian tube sample (Supplementary Fig. 1 and Supplementary Video 1). The resulting fitted cellularities were highly consistent with the *in silico* mixing fractions (Fig. 6a). Simulated cellularities of < 20% could not be fitted to ACN, as already observed in the clinical tissue samples.

**Figure 6.**
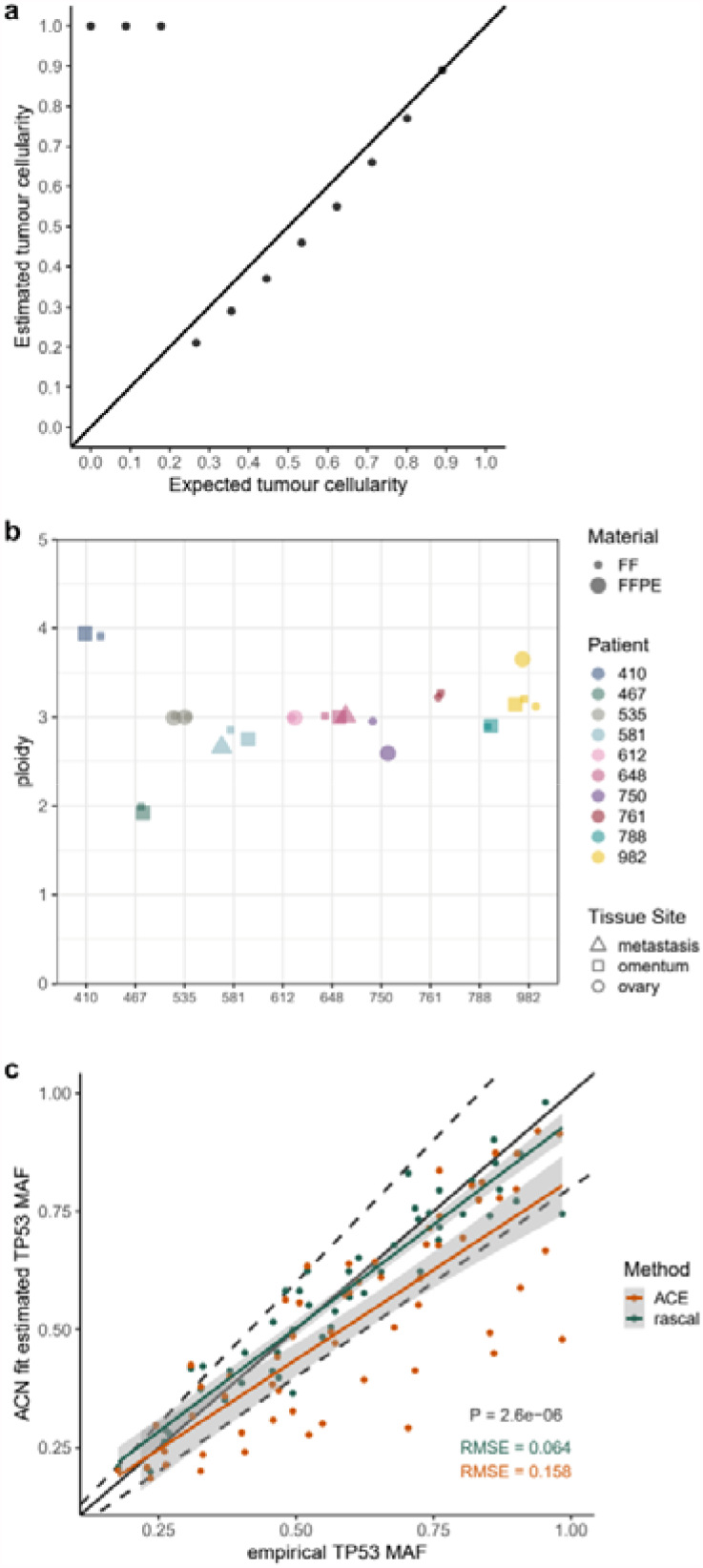
Rascal validation of clinical tissue samples. (**a**) Dilution experiment results for in-silico mixtures of DNA from a high purity tumour tissue and a matched normal fallopian tube tissue at varying proportions. Diagonal line indicates x=y. (see Supplementary Fig. 1 and Supplementary Video 1 for RCN and Supplementary Fig. 6 for ACN profiles). (**b**) Comparison of ploidy estimates obtained from Rascal for HGSOC patients with multi-site or different material tissue samples showing good concordance. Individual patients are shown in different colours, with fresh frozen (FF) vs formalin fixed paraffin embedded (FFPE) samples differentiated by size. Tissue sites are represented as triangles for metastatic samples, squares for samples from the omentum, and circles for ovarian tissue samples. (**c**) *TP53* MAF estimates for HGSOC samples. ACN fits were obtained using Rascal (green) and ACE (orange). Expected *TP53* MAFs were calculated for each ACN fit (y axis) are plotted against empirical *TP53* MAF obtained from TAmSeq. Diagonal line indicates x=y, with dashed lined indicating plus/minus 10% from the diagonal line. RMSE = root median square error. P-value were calculated on the squared errors using the paired one-sided Wilcoxon test.

Secondly, we compared fits achieved for multi-region sampling or fixed and unfixed tissues from the same patient (Fig. 6b). This also showed highly consistent ploidy values across varying samples from the same patient, indicating correct ACN fitting.

Finally, using *TP53* MAF-guided ACN fitting gave significantly more accurate solutions (RMSE = 0.069) compared to unguided fitting using ACE (RMSE = 0.16, P = 2.6 × 10^−6^; Fig. 6c).

This shows that although ACN fitting for high purity samples is easily achievable using Rascal, additional information is required for impure clinical tissue samples. In addition, manual inspection of copy number profiles using Rascal’s interactive web interface allows ACN fitting of otherwise problematic samples.

## Discussion

Studying SCNAs and patterns of genomic instability in cancers is an essential component of pre-clinical and translational science. ACN is required for the accurate identification of actionable driver amplifications in high CIN cancers, which have the poorest outcome in the clinic. Another important benefit of deriving ACN is that it enables inference of copy number signatures^27^, which can identify clinically relevant mutational processes and guide therapy selection. The majority of ACN tools have been developed for deep WGS data. However, for many applications, low-coverage sWGS offers a much more time- and cost-effective alternative to estimate copy number profiles, and is widely accessible in both clinical and research settings.

Here we show that typical and unsupervised use of current tools to analyse low-coverage sWGS data does not yield reliable ACN profiles. This is partly because ACN fitting from sWGS data is very different to ACN fitting from deep WGS; for example, in sWGS we fit total, not allele specific copy number. As a result, many available tools often require more complex input data that cannot be obtained from sWGS and a detailed understanding of how to optimise parameters, making these approaches unsuitable for sWGS-based analyses. This complexity also obscures the understanding of the underlying principles of the algorithms, and prevents basic exploratory data analyses for the reliable interpretation of experimental results.

The Rascal package (relative to <u>a</u>bsolute copy number <u>scal</u>ing) addresses these limitations by providing a significantly improved and validated method and an interactive web application for exploring fits. Using Rascal, robust and accurate ACN estimates can be achieved from sWGS data for sample types of different purities, ranging from cell lines, organoids, ascites and PDX samples to patient samples. For high purity cancer models, Rascal makes ACN fitting highly reliable and easily achievable even for users not trained in bioinformatics.

For clinical tissue samples, ACN fitting is more challenging because both their purity and ploidy is usually unknown. Accurate fits generally require additional information, for example from flow cytometry, FISH^57^, advanced computations histopathology^58^, or MAF estimation of clonal driver genes. We used *TP53* MAFs in HGSOC tumours to illustrate how ACN fits can be achieved from sWGS in clinical tissue samples. In addition, Rascal’s interactive visualisation functionality provides easy inspection of putative ACN fits and is particularly beneficial for the manual curation of low-quality samples, which would have otherwise been discarded by conservative computational tools.

We also show that Rascal is robust to tumour heterogeneity, a general challenge of all bulk sWGS-based approaches, and that resulting ACN fits can indicate the presence of subclonal tumour populations^59^. However, single cell DNA sequencing^60,61^, or deep WGS followed by copy number analyses using tools, such as TITAN^42^ or ASCAT^43^, would be required to study regions of subclonal SCNAs and to estimate the prevalence of clonal clusters^10^.

An obvious limitation of our approach is the current lack of power to obtain ACNs for samples with < 20% cellularity. This could be mitigated by increasing sequencing depth and/or the size of the bin window applied for read counting and copy number segmentation (e.g. from 30kb to 100kb; also see Supplementary Fig. 6). However, further modelling of these sWGS parameters will be required to understand the precise relationship between tumour purity and ploidy and minimum sequencing depths required for accurate ACN fitting^30,35^.

In summary, we provide detailed guidelines and methodological insights for achieving robust and accurate ACN estimates from sWGS data across frequently studied sample types. Rascal and the extensive datasets from this study will provide an invaluable resource for developing and testing new computational approaches for estimating ACN from sWGS.

## Methods

### Cell Lines

Cell lines used in this study are listed in Supplementary Table 3, together with their associated growth conditions and culture media. In general, cell lines were grown according to ATCC/ECACC recommendations. OSE medium is composed of 50:50 medium 199 (Sigma-5017) and medium 105 (Sigma-6395). Cells were tested for mycoplasma contaminations on a regular basis using the qPCR PhoenixDx Mycoplasma kit (Procomcure Biotech), and cell line identities were confirmed prior to DNA extractions using our in-house human short tandem repeat (STR) profiling cell authentication service.

### Clinical samples and primary tissue processing

Solid tumour samples were obtained from patients enrolled in the OV04 study at Addenbrooke’s Hospital, Cambridge, UK. Tumour samples were processed following standardised operating protocols as outlined in the OV04 study design. Tumour samples were cut into small pieces of approximately 0.25 cm^3^ and were subsequently either i) snap-frozen and stored at −80℃ to generate fresh frozen (FF) samples for later DNA extraction, or ii) suspended in freezing media (DMEM:F12 supplemented with 10% DMSO) and stored at −80℃ for surgical implantation into mice for PDX model generation. Additionally, a middle section of the tumour tissue was cut out, suspended in 10% neutral buffered formalin (NBF) for 24 hours and subsequently transferred into 70% ethanol for paraffin embedding and sectioning (FFPE tissue generation).

### Xenograft processing

Tumour tissues from patient-derived xenograft (PDX) bearing mice were processed in a similar way as primary tissue samples outlined above. Tumour bearing mice that reached their endpoint (tumours volumes of no more than 1500 mm^3^) were culled via cervical dislocation or CO_2_ overexposure. Tumours were dissected out and stored in ice-cold PBS until further processing.

As for primary tissues, PDX samples were cut into small pieces and stored for later DNA extraction or PDX implantation work at −80℃. Remaining sample was homogenised using a McIlwain tissue chopper (Brinkmann Vibratome MTC/2E) and subsequently further dissociated in an enzymatic dissociation buffer containing 7 ml DMEM:F12 media, 2.5 ml of 7.5% bovine serum albumin fraction V (Invitrogen, USA), 1 mg ml-1 collagenase A (Roche, UK) and 100 U ml-1 hyaluronidase (Sigma-Aldrich, UK) at 80 rpm for 2 hours at 37℃. Samples were centrifuged at 1500 rpm for 5 min at 4℃ and washed with 12 ml DMEM:F12 followed by another centrifugation step. To further break down clusters of cells, the resulting pellet was suspended in 1 ml of 0.25% trypsin in citrate buffer (STEMCELL Technologies, UK), incubated for 4 min at room temperature, quenched with DMEM:F12, and centrifuged at 1500 rpm for 5 min at 4℃. Subsequently, pellets were resuspended in 1 ml of 5 U/ml dispase (STEMCELL Technologies, UK) and DNase (Sigma-Aldrich, UK) at a final concentration of 0.1 mg/ml for another 4 min at room temperature. Again, cells were washed with DMEM:F12 and spun down. Resulting cell pellets were resuspended in 1 to 2 ml of PBS, depending on cell pellet size, and passed through a 40 µm filter to remove any remaining undigested material. Following filtration, samples were resuspended in PBS making up a final volume of 25 ml. PDX tumour cells were cleared up and isolated from debris, dead cells and blood cells using OptiPrep™ (D1556, Sigma-Aldrich, UK) density gradient centrifugation. Cell samples were assessed for viability and counted using a haemocytometer, and subsequently resuspended in freezing media (10% DMSO in DMEM:F12 + 10% FBS) at 1–10 million cells/vial. Cell aliquots were frozen down at −80°C using “Mr Frosty” containers. Alternatively, for immediate (fresh) injection into mice, single cells were resuspended in injection mix (50 % growth factor reduced Matrigel (BD Bioscience) in PBS).

### PDX passaging

All mouse work conducted was approved and performed within the guidelines of the Home Office UK and the CRUK CI Animal Welfare and Ethics Review Board. Female NOD.Cg-Prkdc^scid^ Il2rg^tm1WjI^/ SzJ (NSG) mice were obtained from Charles River Laboratories. Tumour xenografting was performed either by subcutaneous surgical implantation or subcutaneous injection. For subcutaneous surgical implantation, NSG mice were anaesthetised with isoflurane, treated with analgesics (Carprofen [Rimadyl] at 5 mg/kg), shaved, clipped and disinfected with iodine disinfectant. A small vertical incision of approximately 0.5 cm was then made through the skin over the right flank. A small tumour piece of approximately 2 × 2 × 2 mm from either patient samples or previously established PDXs was suspended in growth factor reduced matrigel and inserted underneath the skin. The incision was closed using surgical glue. For subcutaneous injections, tumour tissue from patients or established PDXs was dissociated as described above and resuspended in injection mix. Following shaving and clipping of NSG mice, approximately 1×10^5^ tumour cells were injected subcutaneously over the right flank in a volume of 50 µl of injection mix. Once PDX tumours reached their endpoint of approximately 1500 mm^3^, tumour tissues were dissected, processed as described above and reimplanted for expansion in serial generations for PDX biobank maintenance.

### DNA extraction

#### 1. Cell Lines

Cell pellets of approximately 1×10^6^ cells were generated from cultured cells for each cell line outlined above and stored at −80℃ until further use. DNA was extracted from cell pellets using the Maxwell® RSC Cultured Cells DNA Kit (Promega, AS1620) with the Maxwell® RSC 48 Instrument (Promega, AS8500).

#### 2. Fresh Frozen tissue samples

Fresh frozen tissue pieces were homogenised using Soft tissue homogenizing CK14 tubes containing 1.4 mm ceramic beads (Bertin) on the Precellys tissue homogenizer instrument (Bertin). Resulting lysates were transferred and subjected to DNA extraction using the AllPrep DNA/RNA Mini Kit (Qiagen) following manufacturer’s recommendations. DNA was eluted in 40 µl Elution buffer.

#### 3. FFPE tissue samples

For each FFPE sample, multiple sections at 10 µm thickness were cut depending on tissue size and tumour cellularity assessed by a pathologist, who marked tumour areas on separate Haematoxylin and Eosin (H&E) stained sections to guide microdissection for DNA extraction. Marked tumour areas from unstained FFPE sections were scraped off manually using a scalpel, dewaxed in xylene and subsequently washed with 100% ethanol. Following complete removal and evaporation of residual ethanol (10 mins at 30℃) DNA was extracted using the AllPrep DNA/RNA FFPE Kit (Qiagen). DNA was eluted in 40 µl Elution buffer.

### H&E purity estimation

H&E sections from FFPE tissues were sent to our pathologist for tumour marking and purity estimation. In addition, H&E sections were scanned and subjected to HALO, an image analysis platform for quantitative tissue analysis in digital pathology. HALO’s random forest classifier was used to separate the H&E image into tumour, stroma and microscope glass slide, allowing tumour purity estimation.

### Shallow Whole Genome Sequencing (sWGS)

DNA extractions were performed as described above, and quantified using Qubit quantification (ThermoFisher, Q328851). DNA samples were diluted to 75 ng in 15 µl of PCR certified water, and sheared by sonication with a target of 200-250bp using the LE220-plus Focused-Ultrasonicator (Covaris) with the following settings: 120 sec at room temperature; 30% duty factor; 180W peak incident power; 50 cycles per burst.

Sequencing libraries were prepared using the SMARTer Thruplex DNA-Seq kit (Takara), with each FFPE sample undergoing 7 PCR cycles, and all other samples undergoing 5 PCR cycles for unique sample indexing and library amplification. AMPure XP beads were used following manufacturer’s recommendations to clean prepared libraries, which were subsequently eluted in 20 µl TE buffer. sWGS libraries were quantified and quality-checked using D5000 genomic DNA ScreenTapes (Agilent 5067-5588) on the Agilent 4200 TapeStation System (G2991AA) before pooling the uniquely indexed samples in equimolar ratios. Pooled libraries were sequenced at low coverage (∼ 0.4 × coverage) on either NovaSeq 6000 S1 flowcells with paired-end 50 bp reads for clinical tissue samples, or the HiSeq 4000 with single 50 bp reads, at the CRUK CI Genomic Core Facility. Resulting sequencing reads were aligned to the 1000 Genomes Project GRCh37-derived reference genome (i.e. hs37d5) using the ‘BWA’ aligner (v.0.07.17) with default parameters.

### Copy number analyses

Relative copy number data was obtained using the QDNAseq^31^ R package (v1.24.0) to count reads within 30, 50 and 100 kb bins, followed by read count correction for sequence mappability and GC content, and copy number segmentation. QDNAseq data were then subjected to downstream analyses using ACE, ichorCNA, ABSOLUTE or our Rascal methodology for ploidy and cellularity estimation and absolute copy number fitting.

#### 1. ACE

ACE absolute copy number fitting was applied to both the cell line and PDX sample sets, as well as our clinical tissue samples, using segmented relative copy number data generated at 30kb bin size with QDNAseq. Absolute copy number data was calculated using the squaremodel function of the ACE BioConductor package with the following parameters based on the author’s recommendations: penalty = 0.5 and penploidy = 0.5. Remaining parameters were set to default values.

#### 2. ichorCNA

The ichorCNA algorithm was applied to both the cell line and PDX sample sets. Prior to ichorCNA, QDNAseq read count data at 50kb bin size was converted into .wig files. Segmentation and prediction of ploidy and cellularity (tumour fraction) were performed using ichorCNA v.0.1.0 (https://github.com/broadinstitute/ichorCNA). Parameters were initialized based on prior knowledge: – normal□=□c(0, 0.05, 0.1, 0.2), –ploidy□=□c(2, 3, 4, 5), with all remaining parameters set to default values.

#### 3. ABSOLUTE

ABSOLUTE was applied to both the cell line and PDX sample sets, using segmented relative copy number data at 30kb bin size obtained from QDNAseq. The RunAbsolute function was used with the following parameters: max.ploidy = 5, max.non.clonal = 0.95, max.neg.clonal = 0.05, primary.disease = “Ovarian Cancer”, platform = “Illumina_WES”, copy_num_type = “total”, min.mut.af = 0.2. All remaining parameters were set to default values. The platform choice (no option available for sWGS) does not impact the analysis as segmented copy number data are provided.

### Rascal - Mathematical framework and underlying principles

Rascal models the copy number output from sWGS data as a function of the ploidy and cellularity of the sample of interest using the same formulation as ACE.

Given a cellularity *c*, a diploid normal genome, and absolute number of tumour copies *a* at a given locus *i*, the average number of copies can be written as

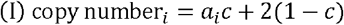

The average copy number across the whole genome in the tumour cells is the ploidy *p*, i.e. the average copy number for a sample made up of tumour and normal diploid cells is

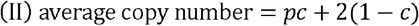

Copy numbers produced by QDNAseq and similar tools are given as ratios rl between the local copy number (equation I) and the average copy number across the genome (equation II) (in some cases, these are log_2_-transformed and referred to as log_2_ ratios):

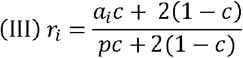

The relative copy number for the zero copy number state, *r*^0^ (*a* = 0), is usually not zero because of the contribution of the normal diploid cells within the sample.

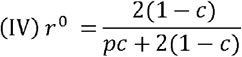

The relative copy number equation can also be rewritten as follows:

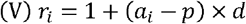

where *d* is the spacing between relative copy numbers for copy number states that differ by one copy, e.g. between the relative copy number observed for a locus with 3 copies and another locus with 2 copies. *d* can be derived from equation (III) given that r_1_ and r_2_ are relative copy number values at locus *i*_1_ and *i*_2_, respectively, in which *a*_2_ −1 = *a*_1_

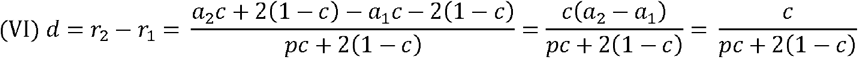

### Rascal – Fitting procedure

The mathematical formulation given in equation (III) can be rearranged to give the absolute copy number (*a*_i_) for each relative copy number value (*r*_i_) for a given ploidy (*p*) and cellularity (*c*):

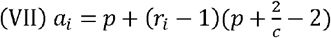

This formulation can be used to perform a grid search on the relative copy number data across various ploidies and cellularities. Rascal assesses the “goodness-of-fit” for each ploidy-cellularity solution by a distance function considering the differences between the scaled values of segmented copy numbers and the nearest whole number for each bin. The distance function is calculated as root mean square deviation (RMSD) or mean absolute deviation (MAD). This provides measures that can be compared across various model fits with differing ploidy and cellularity parameters, with the aim of determining the fit with the lowest RMSD or MAD for each given sample.

Since the RMSD/MAD distance function is solely based on how close scaled copy numbers are to their closest whole integers, this can result in competing best fits. In general, for a given model with ploidy, *p*’, and cellularity, *c*’, there are an infinite number of solutions that have the same spacing, d (see equation (VI)), between successive copy number steps on the relative scale.

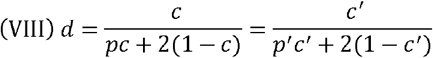

Rearranging this for *c*

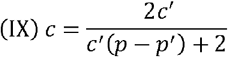

allows us to calculate the cellularity for a higher or lower ploidy solution differing by a whole number (*p* = *p*’ ± 1, 2, 3, …) that will have an equally good fit. This is because shifting the copy number states up or down in whole number increments doesn’t change the proximity of a segment to its closest copy number state. Competing solutions fit the data equally well, thus requiring additional information to identify the most likely ploidy and cellularity.

For pure cancer models, such as cell lines, organoids and PDX samples, best fit solutions were primarily identified based on the lowest MAD distance. Only in cases where multiple solutions (competing fits) with the lowest MAD distance were available, the solution with the highest cellularity was selected.

In clinical tissue samples, where the assumption of high tumour purity cannot be used to guide ACN fitting and distinguish between competing best fits, additional information on either the tumour ploidy or purity, such as SNV allele fractions or an accurate estimate of the tumour cellularity from a histopathologist, is required to obtain reliable ACN fits. We illustrate how this can be achieved using *TP53* mutant allele fractions (MAFs) from Tagged-Amplicon sequencing (TAmSeq), a low-cost ultra-high depth targeted sequencing approach which allows accurate estimation of MAFs, in HGSOC samples:

Rascal ACN solutions are determined as described above. Expected *TP53* MAFs for each putative fit is then calculated using the following formulation in Rascal:

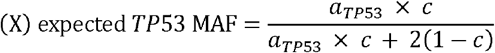

The expected *TP53* MAF for each solution is then compared to the empirical *TP53* MAF (measured by TAmSeq) and the absolute difference is calculated. ACN fits were subsequently ranked based on their MAD and absolute *TP53* MAF difference (MAFdiff), before entering an annotation and solution refinement stage. Samples for which no solution with a matching *TP53* MAF were found (i.e. MAFdiff > 0.3) failed Rascal and were marked for exclusion. Samples for which no empirical *TP53* MAF was available, or which had very low MAFs suggesting low purity (i.e. *TP53* MAF < 0.3) were flagged for manual inspection since low purity samples are more challenging to fit. Manual inspection was then carried out on these samples using the interactive Rascal web application to either confirm or correct for more appropriate ACN fits. Manually curated solutions for flagged samples, and top ranked solutions for all remaining samples were taken forward for downstream analyses.

### Rascal – code availability and accessibility

Rascal is implemented as an R package and can be downloaded from https://github.com/crukci-bioinformatics/rascal. The Rascal interactive web application can be accessed at https://bioinformatics.cruk.cam.ac.uk/rascal.

### Tagged-Amplicon Sequencing (TAmSeq)

Extracted DNA samples were diluted to a final concentration of 10 ng/ml using PCR certified water. Tagged-Amplicon deep sequencing was performed as previously described^47^. In short, libraries were prepared in 48.48 Juno Access Array Integrated Fluidic Circuits chips (Fluidigm, PN 101-1926) on the IFC Controller AX instrument (Fluidigm) using primers designed to assess small indels and single nucleotide variants across the coding region of the *TP53* gene. Following target-specific amplification and unique sample indexing, libraries were pooled and subsequently purified using AMPure XP beads. Quality and quantity of the pooled library were assessed using a D1000 genomic DNA ScreenTape (Agilent 5067-5582) on the Agilent 4200 TapeStation System (G2991AA), before submitting the library for sequencing to the CRUK CI Genomics Core Facility using 150bp paired-end mode on either the NovaSeq 6000 (SP flowcell) or HiSeq 4000 system. Sequencing reads were aligned to the 1000 Genomes Project GRCh37-derived reference genome (i.e. hs37d5) using the ‘BWA-MEM’ aligner. Data analysis and variant calling was performed as previously described^32^.

### Metaphase spreads and chromosome counting

Cells were arrested in metaphase by adding colcemid at a final concentration of 0.075 µg/ml followed by an incubation period of 3 hours. Subsequently, supernatant was removed, cells were collected using trypsin and pelleted. 20 ml of hypotonic solution (50mL UP water,10mL serum and 6mL KCl 0.075M) was added drop-wise to the cell pellet and incubated for 20 minutes. Cells were washed twice with fixative 3:1 (methanol:acetic acid) and once with fixative 3:2. Metaphase preparations were stored in fixative 3:2 at −80℃ until further use. One drop of the metaphase preparation solution was dispensed onto each slide and left to air-dry. Mounting media containing 4′,6-diamidino-2-phenylindole (DAPI) was applied. Metaphase spread slides were imaged on the Operetta CLS imaging system (Perkin Elmer, UK) with the Harmony 4.9 PhenoLOGIC software. Whole slides were scanned first using the 10x objective to identify metaphase spreads, followed by an automated rescan of each spread using the 63x water immersion objective across several z-stacks for chromosome visualisation. Subsequent analysis was performed using the Harmony 4.9 PhenoLOGIC software by segmenting chromosomes within each metaphase spread and the total number of chromosomes per metaphase were counted.

## Author Contributions

C.M.S., M.D.E., F.M. and J.D.B. wrote the manuscript. C.M.S., and M.D.E. performed data analysis. M.D.E. developed the R package and web application. C.M.S., M.V., J.H., and S.B. prepared cell line, PDX and tumour-patient samples and performed experiments. G.M., T.B., F.M. and J.D.B. supervised the work.

## Acknowledgements

We thank all patients who participated in and donated tissue samples to this study. The Addenbrookes Human Research Tissue Bank is supported by the NIHR Cambridge Biomedical Research Centre. We also thank Karen Hosking, Mercedes Jimenez-Linan, and the OV04 study team for their help with clinical tissue samples. We would like to thank the Cancer Research UK Cambridge Institute Genomics, IT & Scientific Computing, Biological Resource Unit, Microscopy, Compliance & Biobanking, and Bioinformatics core facilities for their support with various aspects of this study. We thank Dilrini De Silva and Ania Piskorz for bioinformatics and genomics advice and support. We would like to thank D. Provencher and A.-M Mes-Masson for kindly donating a subset of 25 ovarian cancer cell lines to us that were included in this study. These cell lines were derived with the support of the Banque de tissus et de données of the Réseau de recherche sur le cancer of the Fonds de recherche du Québec - Santé (FRQS) affiliated with the Canadian Tumor Repository Network (CTRNet). Lastly, we would like to thank Bauke Ylstra for critically reviewing the manuscript.

## Extended Data Tables

**Extended Data Table 1.**
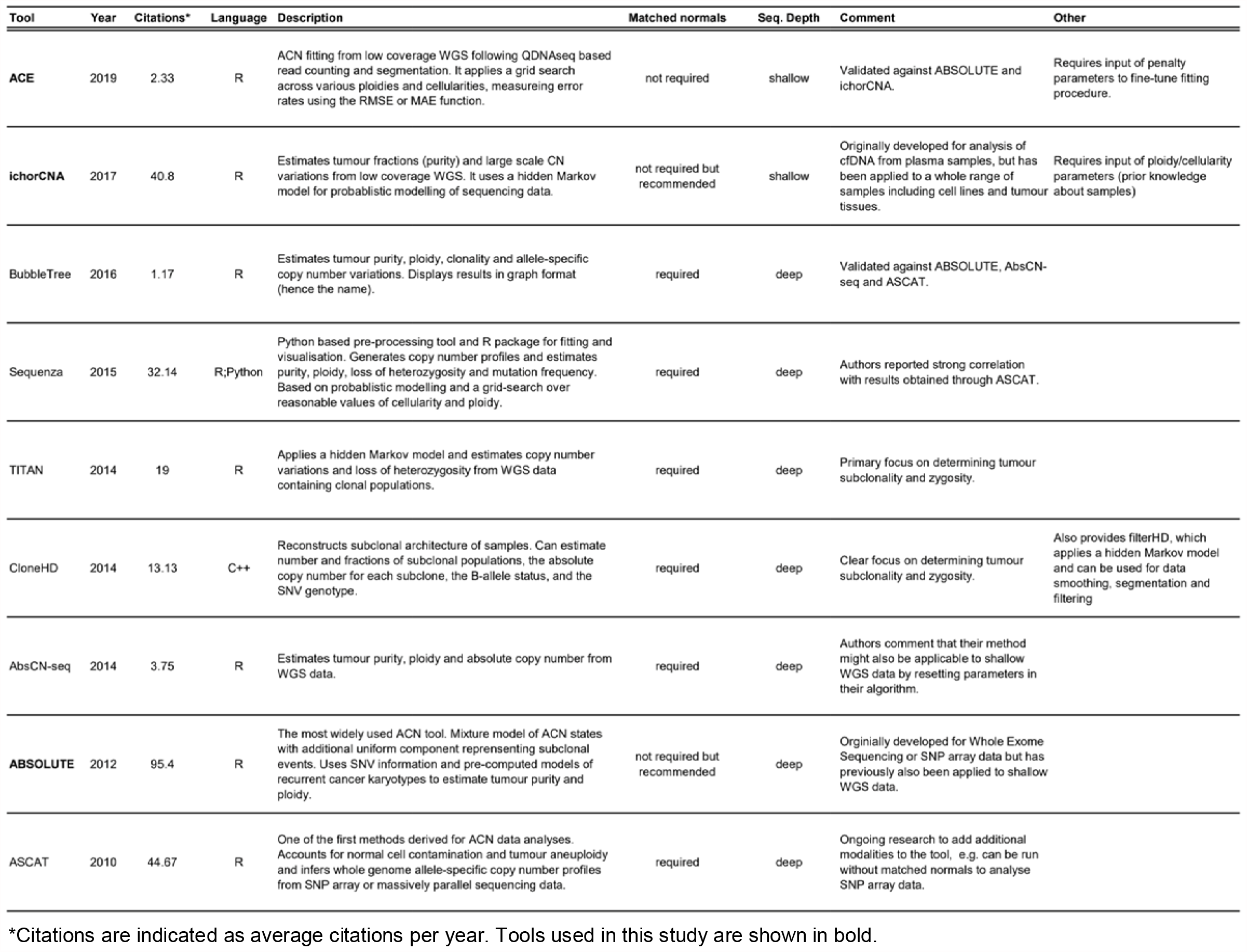
Overview of currently available bioinformatic ACN fitting tools.

## Extended Data Figures

**Extended Data Figure 1.**
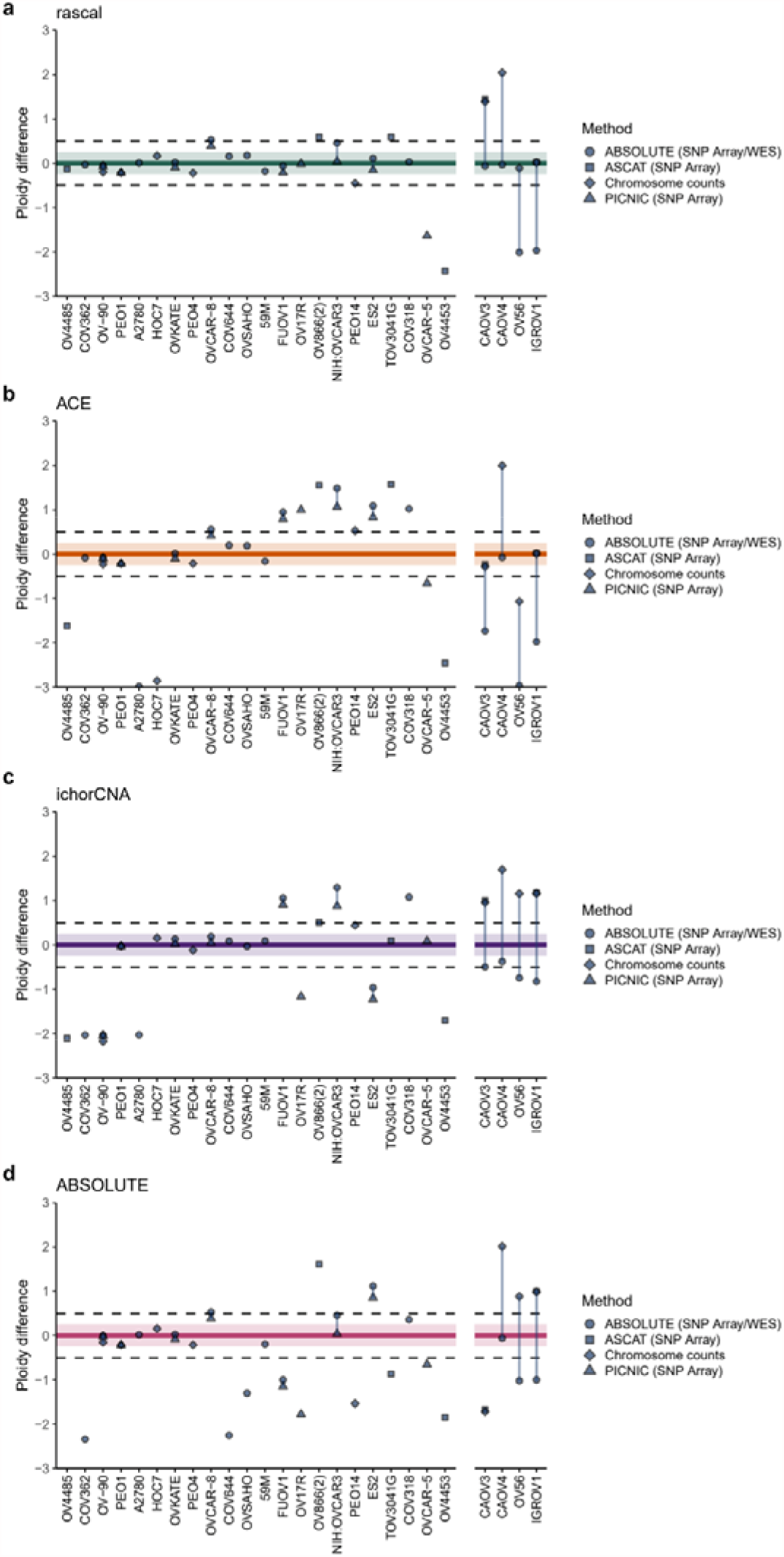
Fitted vs published cell line ploidy across different ACN fitting tools. Fitted ploidies obtained from (**a**) Rascal, (**b**) ACE, (**c**) ichorCNA and (**d**) ABSOLUTE were compared to published ploidy data (See Supplementary Table 1 and 2), where available. (Note that for three of these cell lines assessed via chromosome counting (PEA1, PEA2 and PEO23), no modal chromosome counts (ploidies) could be assigned^52^ for comparison with fitted ploidies and were excluded from downstream analyses.) Data are presented as Bland-Altman plots with the difference between fitted and published ploidies on the y-axis. Ploidy difference of +/−0.25 copy number steps is indicated by shaded ribbons, and ploidy difference of +/−0.5 is indicated by dashed lines. The four cell lines for which discordant published ploidy data was available are shown in the panel to the right. Shapes indicate methods by which published ploidies were determined.

**Extended Data Figure 2.**
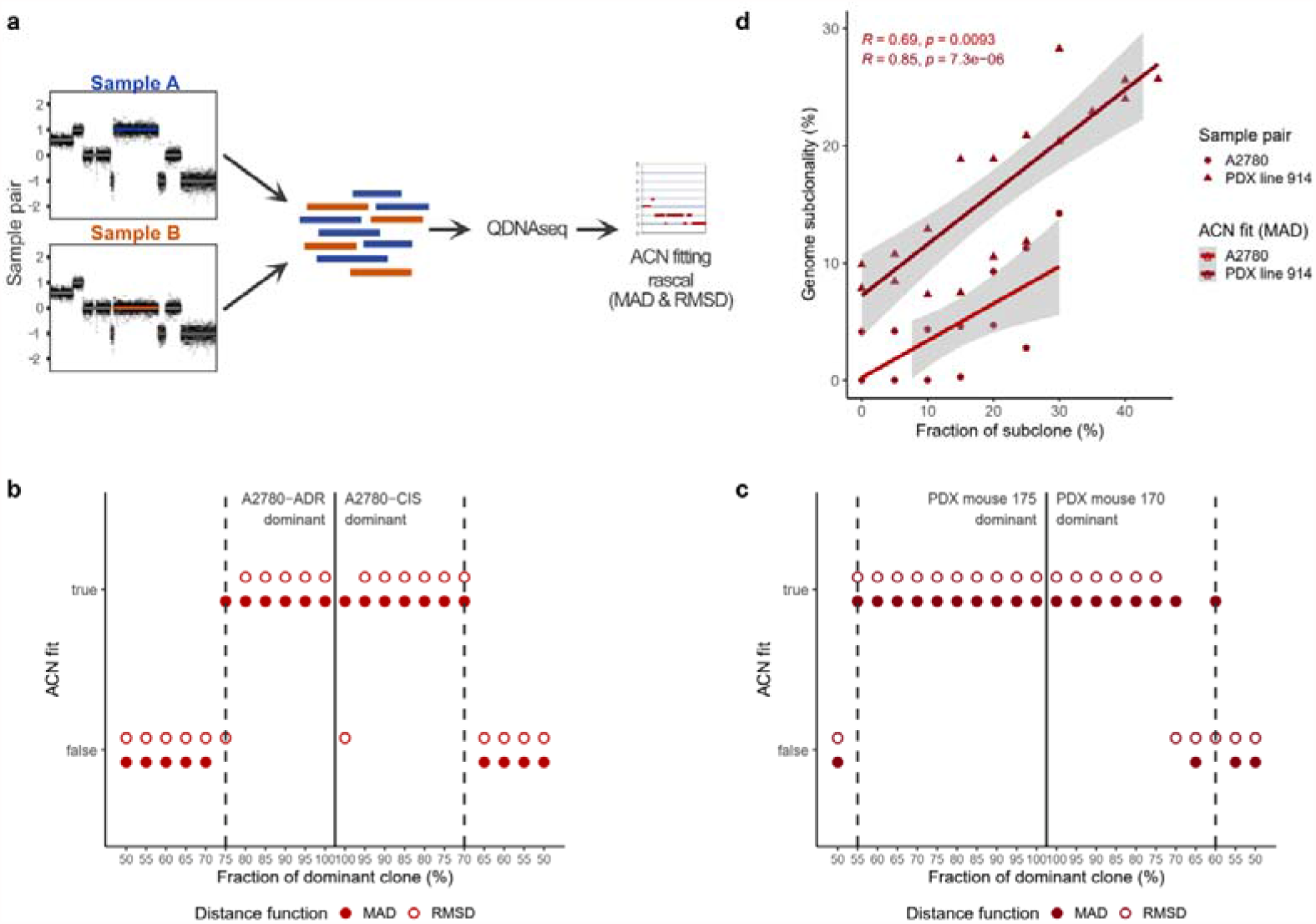
In silico subclonality mixture analysis. (**a**) Schematic of the *in silico* subclonality experiment. QDNAseq read counts from matching sample pairs (e.g. A2780-CIS and A2780-ADR, or PDX line 914 mice 170 and 175) were used to simulate different clonality states with the dominant clone ranging from 50-100% frequency (in 5% increments). Segmented relative copy number data was generated using QDNAseq and applied to Rascal for absolute copy number fitting using both, the MAD and RMSD distance function. Absolute copy number profiles for both sample pairs are shown in Supplementary Fig. 4. (**b**) and (**c**) showing subclonality mixtures derived from the A2780 and PDX line 914 sample pairs, respective, for which correct ACN fits could be obtained using either the MAD (solid circle) or RMSD (hollow circle) distance function. The fraction of the dominant clone for each admixture is shown along the x-axis. The dashed, vertical, grey lines indicate the minimum dominant clone fraction at which correct ACN fits could be obtained. (**d**) Percentage of genome subclonality calculated as before (Fig. 4e) compared to the *in silico* simulated subclone fraction. Only samples which were fitted to their correct ACN fits using the MAD distance function were included. A2780 derived samples are shown as solid circles; PDX line 914 derived samples as triangles.

