## Supplementary Figures 1-6 for "Absolute copy number fitting from shallow whole genome sequencing data"

#### Contents in this document

### Other Supplementary Material

(provided separately)

Tables:

1. Supplementary Table 1 – Cell line ACN fits obtained from ABSOLUTE, ichorCNA, ACE and Rascal.
2. Supplementary Table 2 – Available published ploidy data for subset of 30 cell lines.
3. Supplementary Table 3 – Cell line growth conditions and information

Videos:

1. Supplementary Video 1 – The impact of tumour purity on relative copy number signal
2. Supplementary Video 2 – Example: PDX patient line 914 (Rascal fits)
3. Supplementary Video 3 – Example: PDX patient line 716 (ACE fits)

***Supplementary Figure 1 – Effect of tumour purity on relative copy number signal***

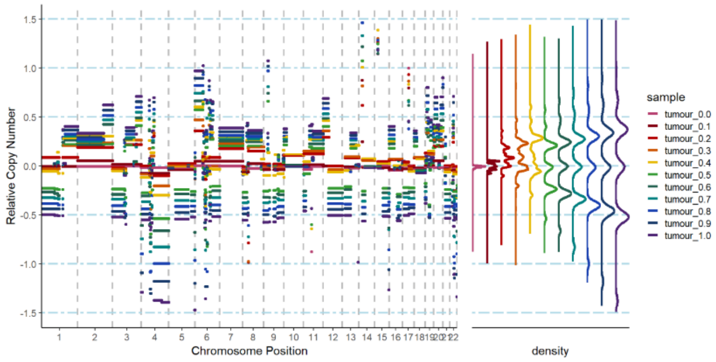

**Relative copy number signal decreases with decreasing tumour cellularity**. Dilution experiments were performed by *in-silico* mixing reads from a high purity tumour tissue (*TP53* MAF ≅ 0.94) with reads obtained from a matched normal fallopian tube sample. Relative copy number segments are plotted with different colours indicating different mixtures. Copy number segment density curves are shown on the right panel of the plot for each mixture using the same colour coding.

***Supplementary Figure 2 – TP53 based tumour purity estimation of PDX tissue samples***

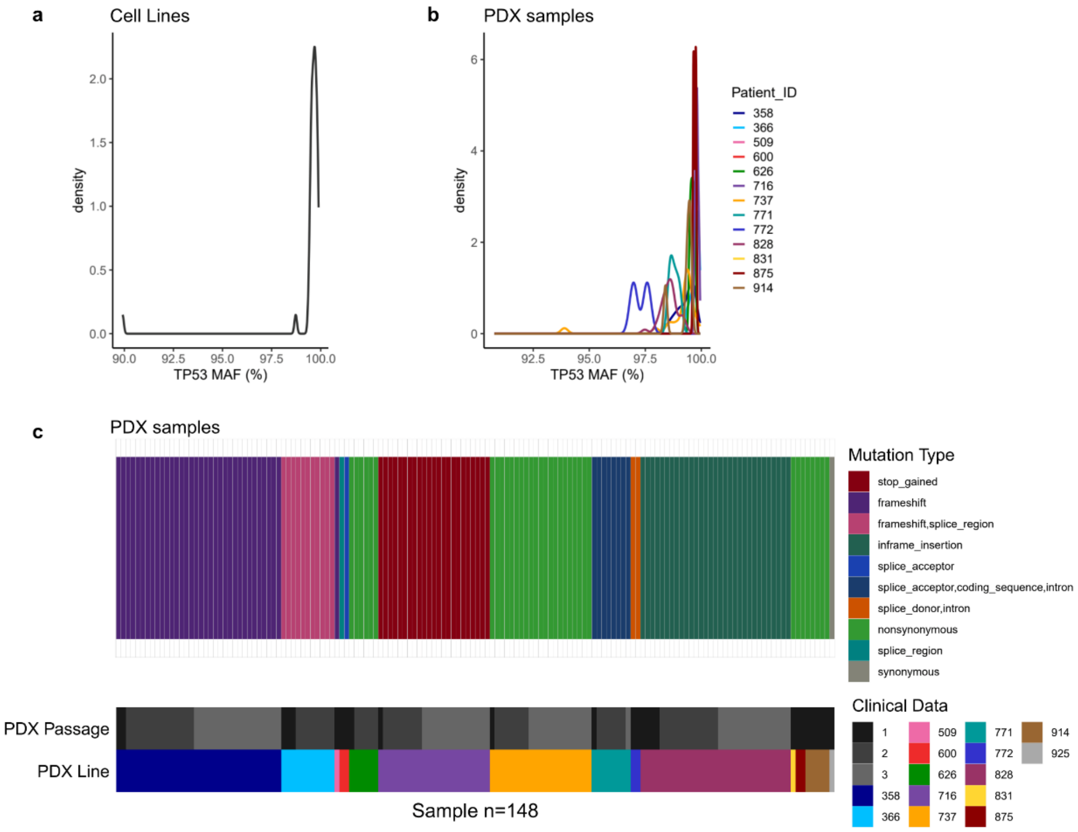

(**a**) Density distribution of *TP53* mutant allele fractions (MAFs) for cell line samples. Note that only cell lines of the HGSOC subtype are plotted (n = 41) where available. (**b**) Density distribution of *TP53* MAFs for PDX samples grouped and coloured by PDX line (Patient ID). (**c**) Oncoprint plot showing detected *TP53* mutations for PDX tissue samples are conserved between samples and PDX passages from the same patient line. Samples are grouped and ordered by PDX line (Patient ID) and passage number. Mutation Types are indicated by different colours.

***Supplementary Figure 3 – ACN fitting MAD vs RMSD error function on CAOV4 cell line***

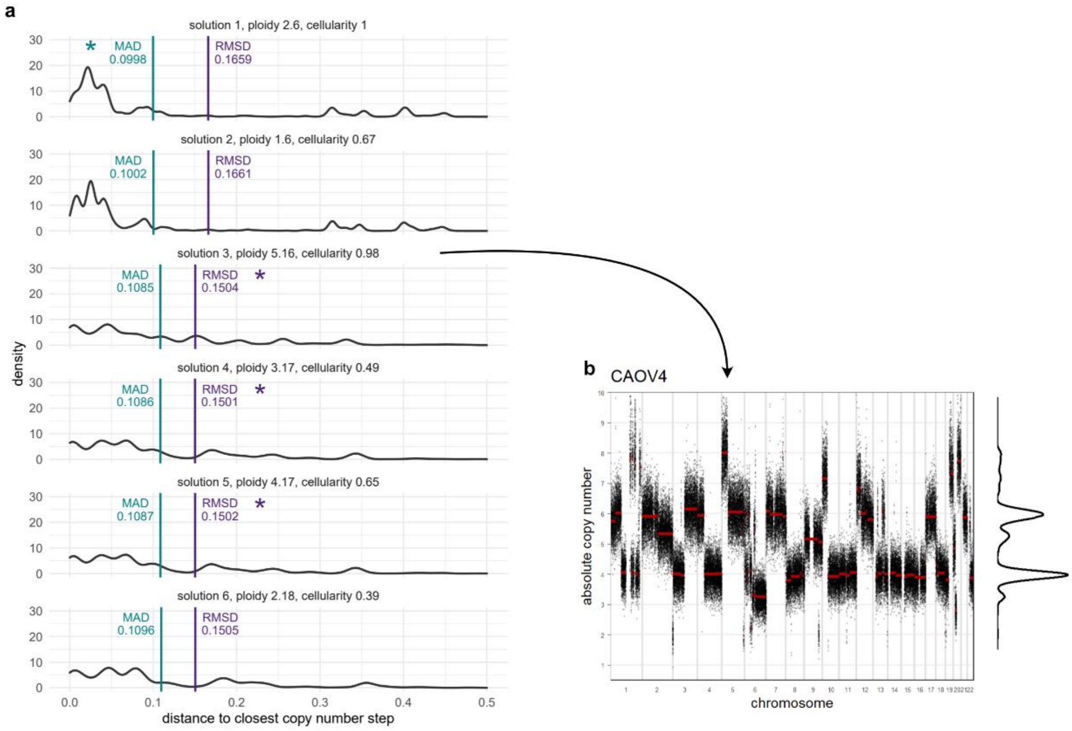

(**a**) Top six ACN solutions ordered by MAD values (lowest to highest). Solutions favoured by MAD and RMSD distance functions are indicated by teal and purple asterisk, respectively. (**b**) Incorrectly fitted absolute copy number plot according to top RMSD-favoured solution with cellularity of 0.98, and ploidy 5.16, overcompensating for intermediate segments arising from subclonal populations.

***Supplementary Figure 4 – In silico subclonality mixture ACN profiles***

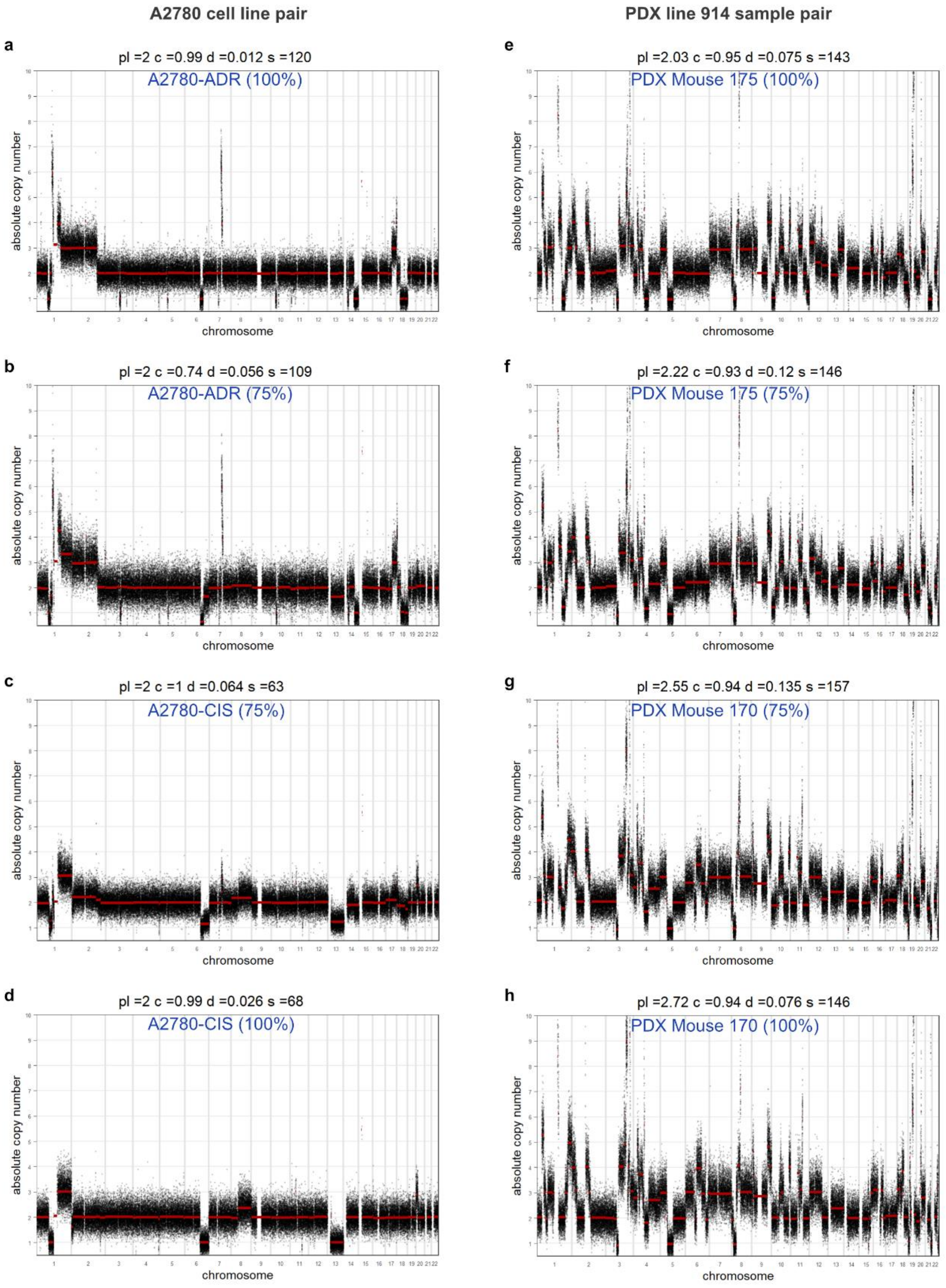

Absolute copy number profiles for A2780 (left column) and PDX line 914 (right column) sample pairs. (**a**) 100% A2780-ADR, (**b**) 100% A2780-ADR, (**c**) 75% A2780-CIS, and (**d**) 100% A2780-CIS. (**e**) 100% PDX Mouse 175, (**f**) 75% PDX Mouse 175, (**g**) 75% PDX Mouse 170, and (**h**) 100% PDX Mouse 170. (pl = ploidy; c = cellularity; d = distance measure (MAD); s = number of segments).

***Supplementary Figure 5 – Competing best fit solutions***

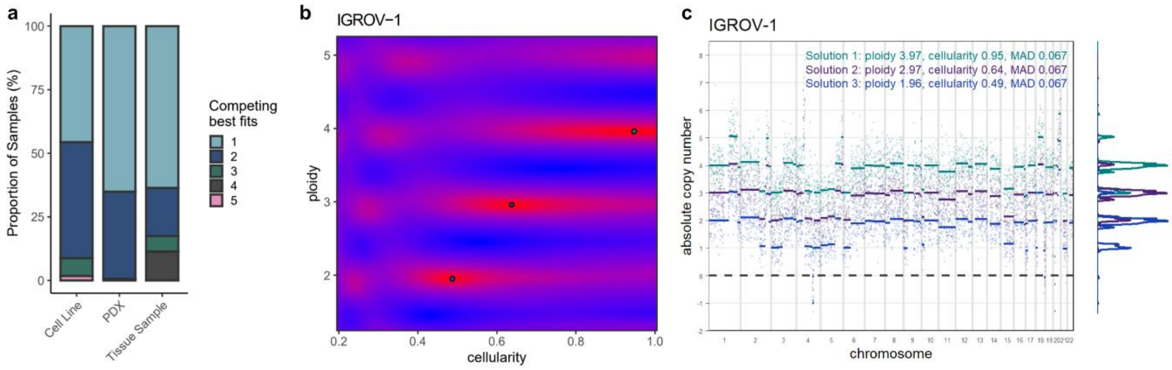

(**a**) Number of competing best fits (based on the MAD distance function) for cell lines, PDXs and clinical tissue samples. An example of a cell line for which Rascal generates three competing solutions that fit the data equally well is given using IGROV-1. (**b**) Heatmap representing the ACN solution grid search for IGROV-1. Cold areas (poor fits) are shown in blue, and hot areas (good fits) are shown in red. The best solutions are indicated by dots. (**c**) Absolute copy number profiles for each of the three competing solutions. Solution 1 (ploidy 3.97; cellularity 0.95), solution 2 (ploidy 2.97; cellularity 0.64) and solution 3 (ploidy 1.96; cellularity 0.49) are shown in teal, purple and blue, respectively. Density plots indicating the distribution of copy number segments across different copy number states are shown to the right of the plot for each solution. Using the assumption of high purity, we can infer the first solution (ploidy 3.97, cellularity 0.95) to be correct; and indeed IGROV-1 is tetraploid[48,50]. This example emphasizes the importance of prior knowledge to distinguish between competing solutions[46], and highlights the challenges of ACN fitting in clinical tumour samples.

***Supplementary Figure 6 – ACN fitting at 30 and 100kb across different tumour purities***

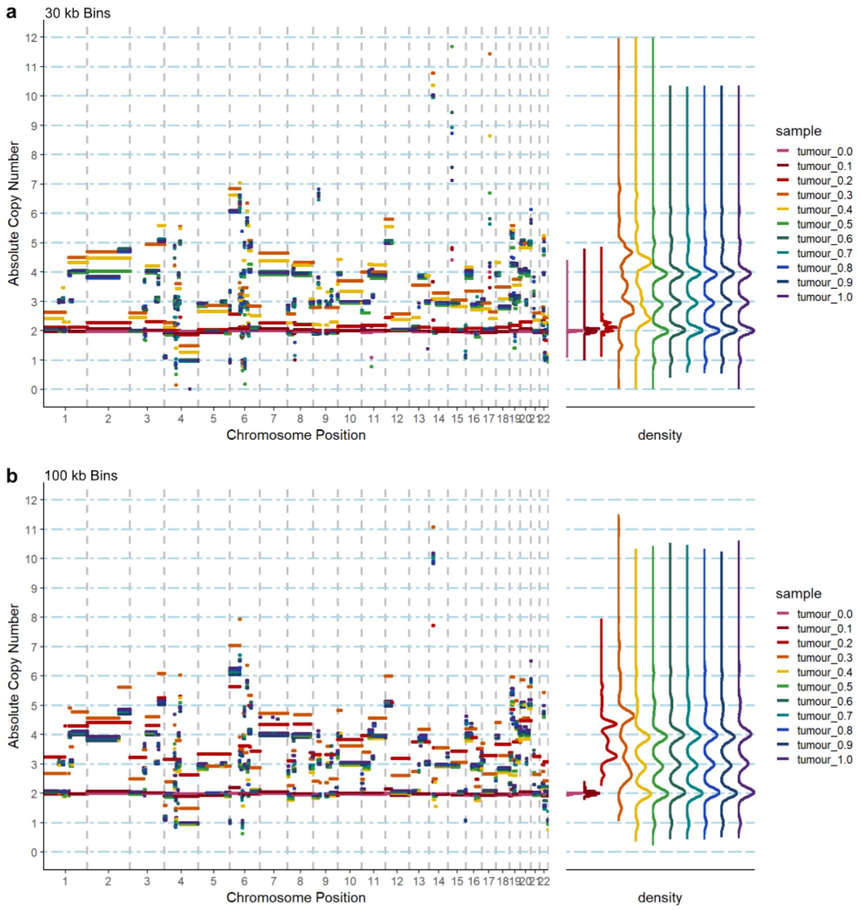

Absolute copy number fitting performed on dilution experiment data (See Supplementary Figure 1 for relative copy number data). Absolute copy number segments are plotted with different colours indicating the different dilution mixtures for (**a**) 30 kb, and (**b**) 100 kb bin sizes. Copy number segment density curves are shown to the right of each plot.
