## Supplementary figures and images for "Absolute copy number fitting from shallow whole genome sequencing data"

### Supplementary Video 1

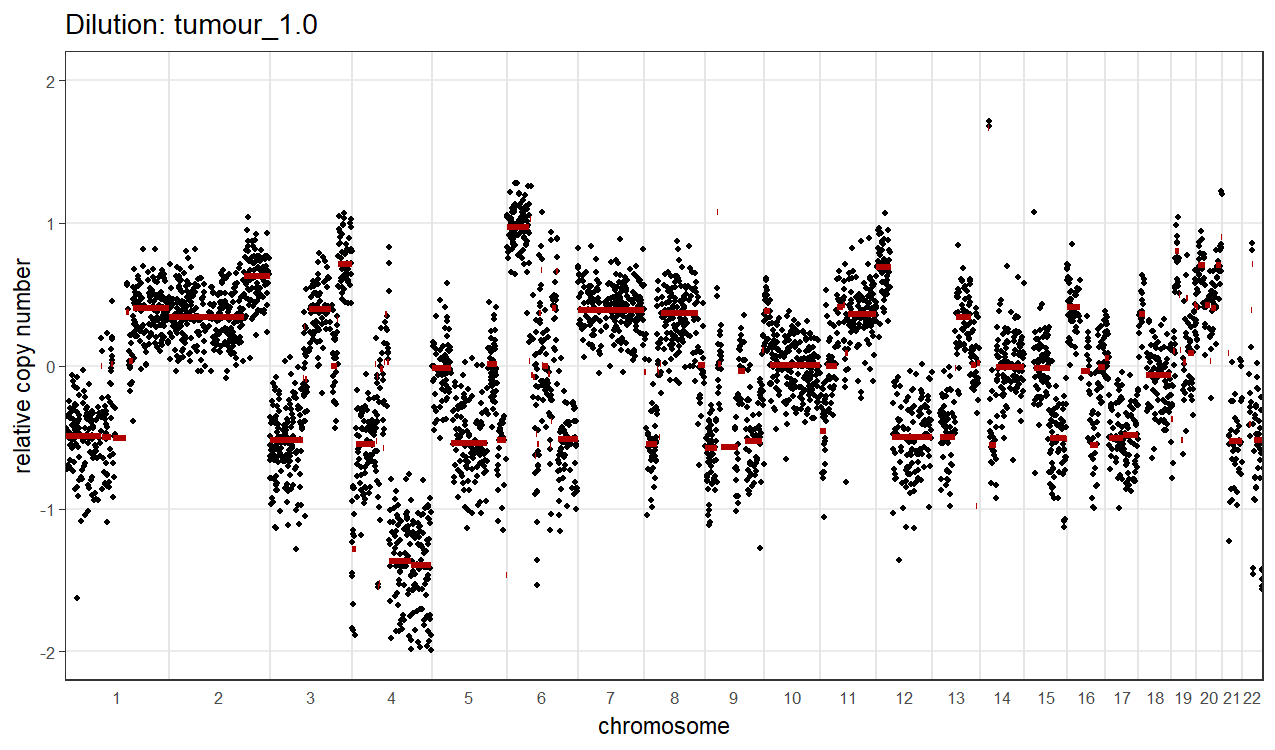

### Supplementary Video 2

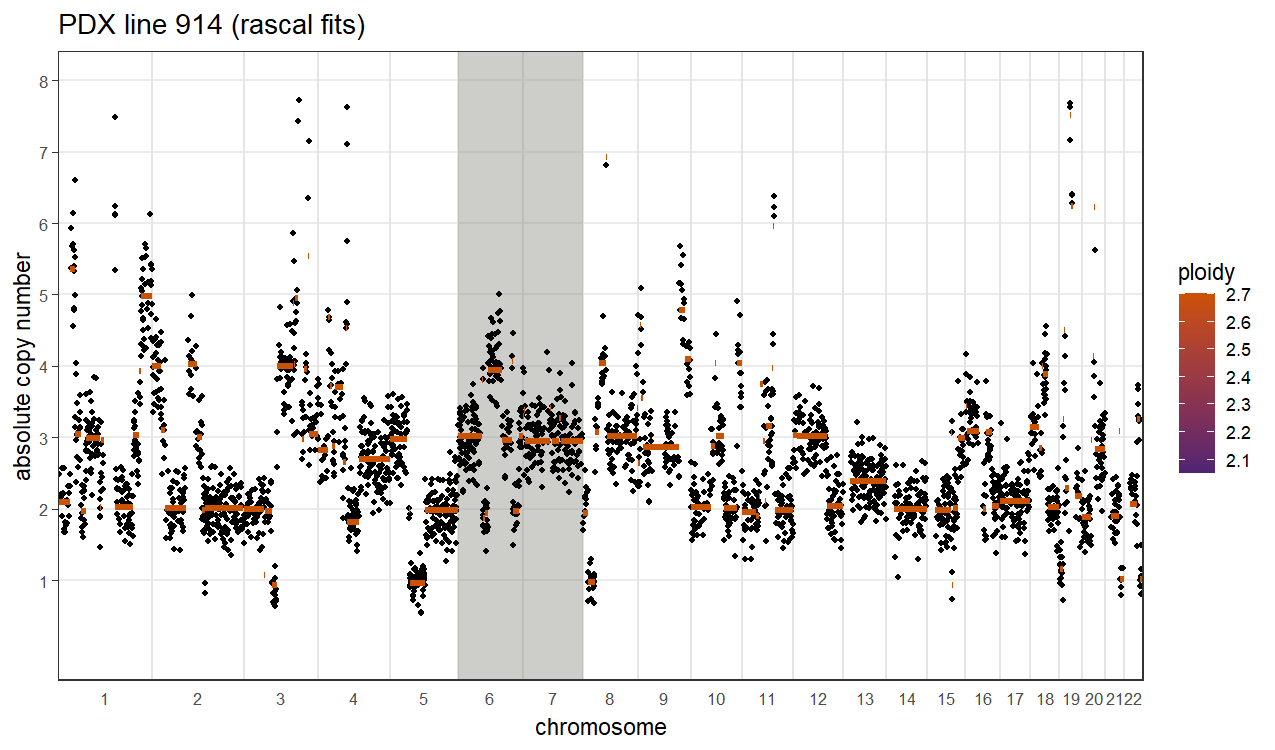

### Supplementary Video 3

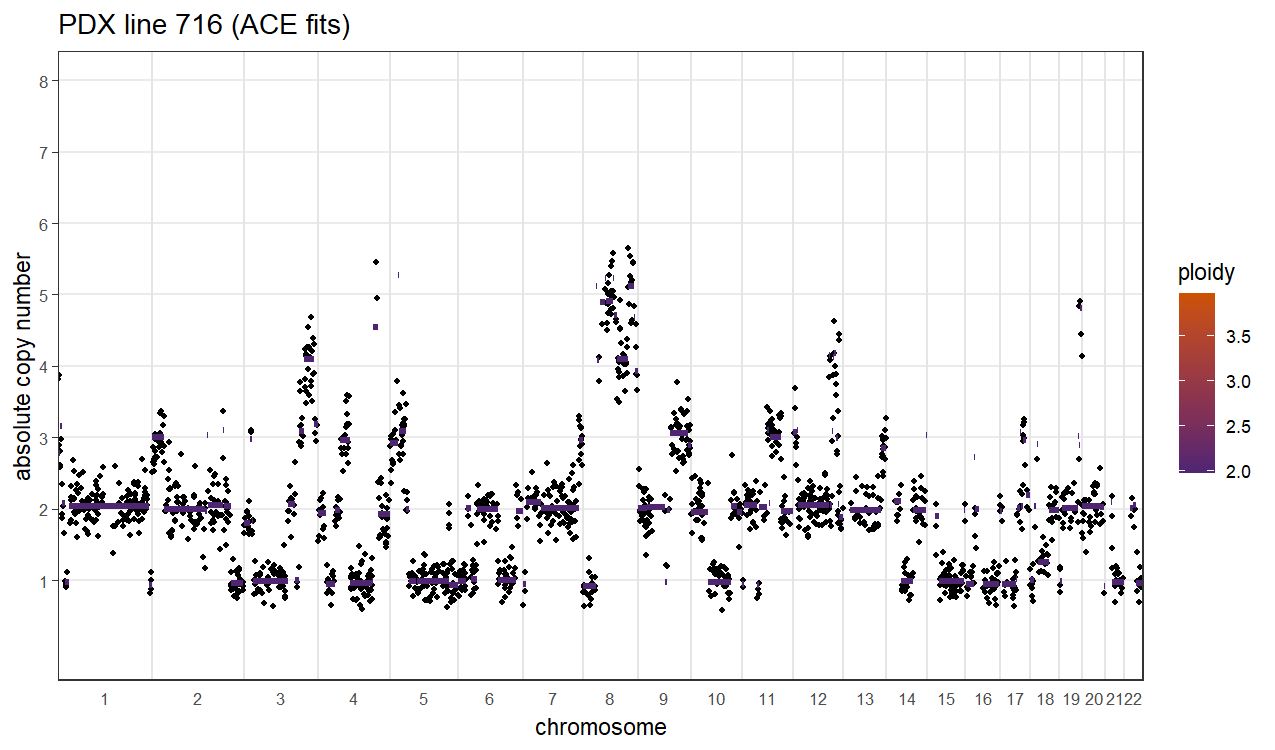
